# Structural basis for regulation of CELSR1 by a compact module in its extracellular region

**DOI:** 10.1101/2024.01.26.577439

**Authors:** Sumit J. Bandekar, Krassimira Garbett, Szymon P. Kordon, Ethan Dintzner, Jingxian Li, Tanner Shearer, Richard C. Sando, Demet Araç

## Abstract

The Cadherin EGF Laminin G seven-pass G-type receptor subfamily (CELSR/ADGRC) is one of the most conserved among adhesion G protein-coupled receptors and is essential for animal development. The extracellular regions (ECRs) of CELSRs are large with 23 adhesion domains. However, molecular insight into CELSR function is sparsely available. Here, we report the 3.8 Å cryo-EM reconstruction of the mouse CELSR1 ECR and reveal that 14 domains form a compact module mediated by conserved interactions majorly between the CADH9 and C-terminal GAIN domains. In the presence of Ca^2+^, the CELSR1 ECR forms a dimer species mediated by the cadherin repeats putatively in an antiparallel fashion. Cell-based assays reveal the N-terminal CADH1-8 repeat is required for cell-cell adhesion and the C-terminal CAHD9-GAIN compact module can regulate cellular adhesion. Our work provides molecular insight into how one of the largest GPCRs uses defined structural modules to regulate receptor function.

## Introduction

Multicellular organisms use an array of cell-surface receptors to facilitate productive adhesion between cells and initiate events to control developmental and regulatory processes. The adhesion class of G protein-coupled receptors (aGPCRs) are an understudied family of cell-surface receptors that link adhesion to intracellular events through their multidomain extracellular regions (ECRs)^1–3^. The aGPCR subfamily of Cadherin EGF Laminin-G seven-pass G-type receptors (CELSRs or ADGRCs) are one of two aGPCR subfamilies conserved as distantly as *C. elegans* and CELSRs are essential for the embryonic and neural development of animals^4–7^. There are three mammalian CELSRs (CELSR1-3) and other commonly studied orthologs include *C. elegans* Flamingo (Fmi) and *D. melanogaster* Starry Night (Stan)/Flamingo (Fmi)^8^. CELSRs are involved in the process of planar cell polarity (PCP)^9–14^, where they are key for neural tube closure^9,11,15,16^, organization of inner ear stereocilia^9,11^ and neuroepithelial cilia^17^, lymphatic valve formation^18^, and skin hair patterning^9,16^. In the nervous system, CELSRs regulate neuronal migration, dendritic growth, axon guidance, and glutamatergic synapse formation^19–24^. CELSRs are ubiquitously expressed in epithelial cell types during embryonic development^25–28^. During development of the nervous system, CELSRs are highly expressed throughout the CNS^25–27^. Mutations in CELSRs are strongly associated with neural tube defects including spina bifida and craniorachischisis, resulting in a range of physical and intellectual disabilities up to embryonic lethality^29–33^. CELSRs are also associated with lymphedema, Joubert syndrome, Tourette syndrome, fetal hydrops, and the progression of leukemia^34–42^.

From their N-termini, CELSRs have poorly conserved prodomains (PRO) with a putative furin cleavage site. Following PRO are nine cadherin repeats (CADH1-9) which take up ∼ 40 % of the ECR mass, making CELSRs a unique CADH-containing subfamily of the aGPCRs^43–45^. Cadherin repeats are protein-protein interaction modules with ∼ 100 residue globular domains with linker regions that coordinate calcium. Three Ca^2+^ ions bind between two CADH repeats and rigidify this linker and this can include many repeats in tandem. This rigid CADH repeat structure mediates cell adhesion in a specific manner^46,47^. CADH repeats act as ‘molecular Velcro’ to hold cells together and are mechanotransducers^48^. CELSR1 mutants which change single CADH residues directly involved in coordinating Ca^2+^ result in PCP phenotypes and embryonic lethality^9^. Following the CELSR cadherin repeats, there is a Flamingo box (Fbox) domain which is unique to CELSR/Fmi proteins^14^. Fbox domains bear structural homology to the membrane-adjacent domains (MADs) of protocadherins (pCDHs)^49^. Furthermore, CELSRs have a similar domain architecture to pCDHs and FAT cadherins, which have a series of CADH repeats, a MAD domain, and either a transmembrane domain (pCDH) or a series of LamG and EGF domains (FAT)^43,45,49–51^. pCDHs overlap several N-terminal CADH repeats to dimerize, making tight antiparallel dimers which can serve as mechanotransducers; these proteins are also involved in neuronal self-avoidance^52–57^. MADs can form cis-dimers with an adjacent CADH repeat where the minimal unit of N-CADH-MAD-C is necessary to dimerize^49^. Multiple studies have demonstrated the importance of the CELSR CADH region in cell adhesion, but molecular detail is sparsely available^12–14,21^. The only available information comes from a recent structural study, which determined low resolution images of the full extracellular region of human CELSR2, showing an extended region likely consisting of the CADH repeats, and a globular region of unknown structure. This study also shows that the CADH repeats likely wrap around each other tightly, using multiple cadherin domains to form tight extended dimer species similar in principle to protocadherins^58^. Another recent study determined crystal structures of monomeric CELSR1 CADH1-4 and CADH4-7, giving the first high-resolution insight into the CADH repeat region of CELSR and revealing a non-canonical linker between CADH5 and CADH6^59^.

Next, CELSRs have a series of epidermal growth factor (EGF) and Laminin G (LamG) domains. CELSR1 has 3 EGF repeats (EGF1-3), the first LamG domain (LamG1), EGF4, LamG2, and EGF5-8. EGF repeats found extracellularly can serve scaffolding functions^60–64^ or act as protein-protein interaction (PPI) modules^65^. LamG domains are found in PPI sites such as in Neurexin^66,67^. The ECR ends with the hormone receptor (HormR) and GPCR autoproteolysis inducing domains (GAIN). CELSR orthologs across different species variably include the residues for GAIN domain autoproteolysis^68,69^.

In *C. elegans* Fmi, the EGF/LamG/HormR/GAIN region plays a role in the interaction with Frizzled, an Fmi binding partner involved in PCP^70^, but no function for this region has been demonstrated for vertebrate CELSRs. At their C-terminus, CELSRs have a seven-transmembrane (7TM) region and a long intracellular region of about 300 residues.

Despite various hints of CELSR structure and function available in the literature, detailed information at the molecular level about these proteins is minimally present. In addition, mechanistic insight for how CELSRs receive adhesion as an extracellular cue and process it remains unavailable. In this work, we determined the 3.8 Å cryogenic electron microscopy (cryo-EM) reconstruction of the CELSR1 ECR, which revealed a compact multidomain module (CMM) consisting of fourteen domains including CADH9-GAIN. Within this CMM, the N-terminal CADH9 and C-terminal GAIN domain form the primary contact interface using conserved residues. We also report the experimental structure of the Fbox domain within this module. We used biophysical assays to study the full ECR in the presence of Ca^2+^ and we observed an extended species, which we propose represents an intertwined antiparallel dimer found in a configuration similar to protocadherins. We analyzed truncations of CELSR1, and we show the CADH1-8 module is sufficient for cellular adhesion. Finally, we show that point mutations in the CADH9/GAIN interface led to increased cell-aggregation activity. Other aGPCRs have CMMs and therefore the CMM may represent a paradigm for regulating aGPCR function which maintains specific distance restraints, allows regulation by splice variation, or is sensitive to mechanical force.

## Results

### Cryo-EM reveals that CADH9-GAIN forms a compact module

We purified the CELSR1 ECR construct containing domains from CADH1-GAIN (Fig. 1a, Supplementary Fig. 1a-c) and we performed single-particle cryo-EM analysis on this construct. Our initial 2D class averages from screening images showed compact triangle-shaped particles about 160 Å in the longest dimension (Supplementary Fig. 1d, e). Processing of a full dataset resulted in a 3.8 Å reconstruction (Fig. 1b, Table 1, and Supplementary Fig. 2, Supplementary Fig. 3) which allowed for unambiguous assignment of the CADH9-GAIN domains within the density map (Fig. 1c). The CADH1- 8 domains are unresolved in this reconstruction; presumably they are flexible relative to the CADH9-GAIN module in the absence of Ca^2+^. They appear in 2D averages as blurry density connected to CADH9 (Supplementary Fig. 2c). The fourteen domains in the CADH9-GAIN module form a compact multidomain module (CMM) architecture reminiscent of the ouroboros, an ancient symbol of rebirth where a snake is depicted eating its own tail^71^. The overall dimensions are ∼ 160 x 120 x 75 Å. CADH9 forms the ‘head’ of the snake (Fig. 1c), and the GAIN domain is the ‘tail’. The CADH9/GAIN interaction surface plays a critical role in stabilizing the module (Fig. 1c). CADH9, Fbox, EGF1-3, and LamG1 are tightly packed next to each other, occluding the putative LamG1 binding site^72^. EGF4 and LamG2 also form a compact group with the LamG2 putative ligand binding site^72^ accessible for binding partners. Then, EGF5-8 are found in an extended conformation, representing the long body of the snake wrapped around the other domains. EGF6 contacts LamG1, and EGF7 contacts EGF2. Finally, EGF8 leads into the HormR domain and the GAIN domain.

**Fig. 1.**
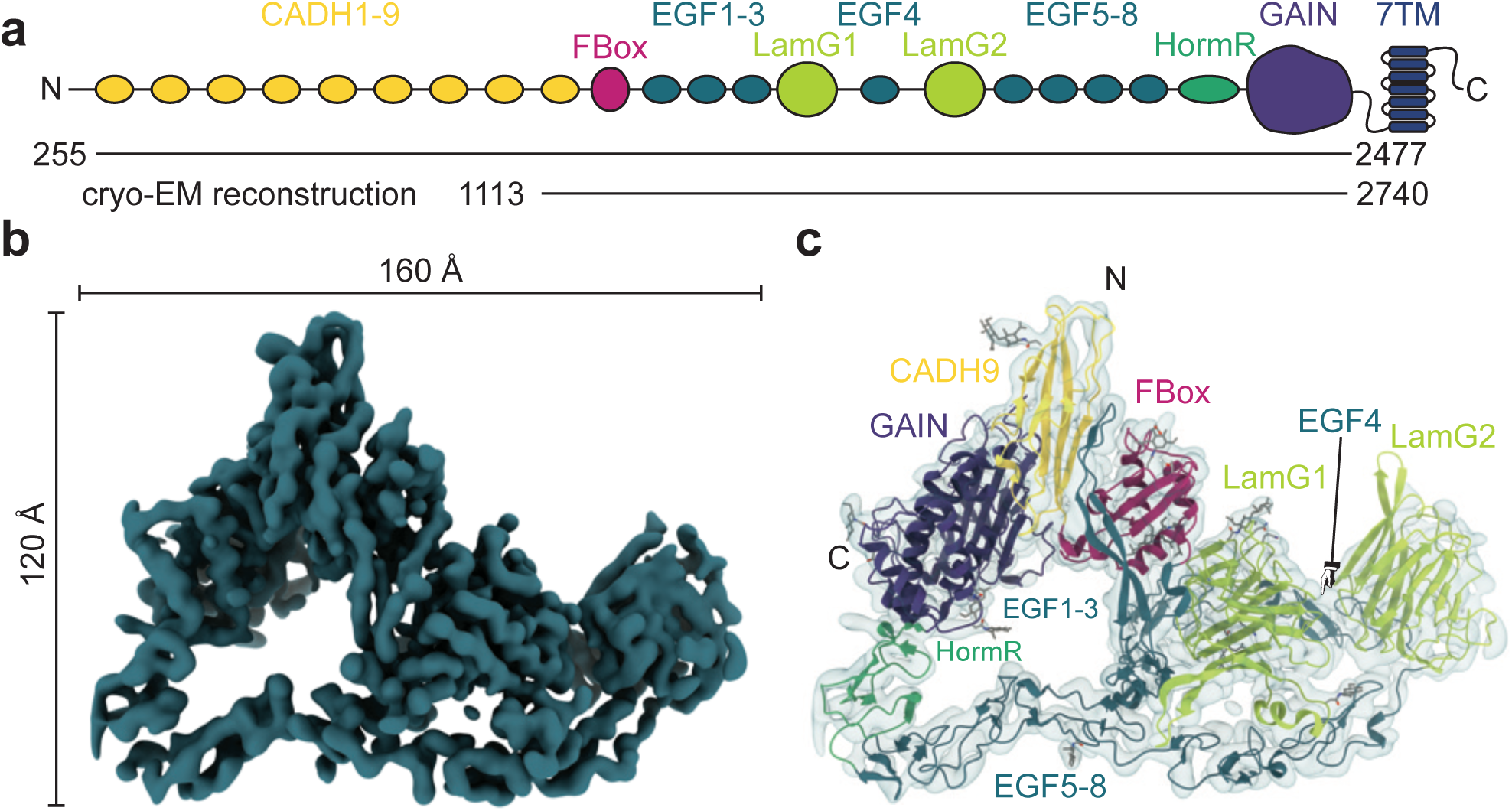
The 3.8 Å Cryo-EM reconstruction of the extracellular region of CELSR1 reveals a compact interdomain bundle including 14 domains. **a** A cartoon diagram shows the overall domain architecture of CELSR1 with horizontal lines indicating boundaries used in the construct and the boundaries resolved in the reconstruction. PRO: prodomain; CADH: cadherin repeat; Fbox: Flamingo Box domain; EGF: epidermal growth factor repeat; LamG: laminin G repeat; HormR: hormone receptor domain; GAIN: GPCR autoproteolysis-inducing domain; 7TM: seven transmembrane region; ICR: intracellular region. **b** Coulomb potential map of the cryo-EM reconstruction reveals a compact triangular shape. **c** A cartoon representation of the CELSR1 ECR atomic model colored as in **a** fit into the coulomb potential map shows that the N-terminal portion of the structure interacts with the C-terminal portion.

**Table 1.** Cryo-EM Data Collection and Processing Statistics.

|  | CELSR1 ECR Dataset A | CELSR1 ECR Dataset B |
| --- | --- | --- |
| ***Data Collection and Processing*** |  |  |
| Magnification (x) | 81,000 | 81,000 |
| Voltage (kV) | 300 | 300 |
| Electron Exposure (e<sup>-</sup>/Å<sup>2</sup>) | 50 | 50 |
| Exposures | 5826 | 5550 |
| Defocus range (μm) | -1.0 to -2.0 | -0.7 to -1.7 |
| Pixel Size (Å) | 0.55 | 0.55 |
| Symmetry Imposed | C1 | C1 |
| ***Reconstruction*** |  |  |
| Initial Particle images | 4,964,141 |  |
| Final Particle images | 275,998 |  |
| Global Resolution (Å) | 3.8 |  |
| Resolution Range (Å) | 2.4 – 6.0 |  |
| Map Sharpening B factor (Å<sup>2</sup>) | -193 |  |
| ***Model Composition*** |  |  |
| Non-hydrogen atoms | 10374 |  |
| Protein Residues | 1319 |  |
| Ligands | 15 |  |
| ***B Factors*** |  |  |
| Protein (Å<sup>2</sup>) | 270 |  |
| Ligand (Å<sup>2</sup>) | 271 |  |
| ***R.M.S. deviations*** |  |  |
| Bond lengths (Å) | 0.005 |  |
| Bond Angles ( ° ) | 0.908 |  |
| ***Validation*** |  |  |
| MolProbity Score | 2.11 |  |
| Clashscore | 13 |  |
| Poor rotamers (%) | 0.97 |  |
| ***Ramachandran Plot*** |  |  |
| Favored (%) | 91.94 |  |
| Allowed (%) | 7.6 |  |
| Disallowed (%) | 0.46 |  |

Our cryo-EM reconstruction allows us to report the experimental structure of the Fbox domain, a domain type thought to be unique to CELSR/Flamingo orthologs (Supplementary Fig. 4a)^14^. We confirm previous work which predicted the Fbox domain to be structurally similar to MAD domains of protocadherins and FAT cadherins as well as the ferredoxin-like domains of nuclear pore proteins^49,73^. Using the DALI webserver, we identified several SEA domains as structural homologs of Fbox, including SEA of the aGPCR ADGRG6 (Supplementary Fig. 4a)^74^. The CELSR Fbox domain does not have the residues for furin cleavage which some SEA domains have (Supplementary Fig. 4b)^75^. The pCDH15 MAD domain forms a dimer with its adjacent CADH repeat, where the CADH of one monomer interacts with the MAD of the other monomer, and vice versa. The orientation of CAHD9 to Fbox is different from the CAHD11-MAD orientation in pCDH15, and the dimerization surface is sterically hindered within the CELSR CMM (Supplementary Fig. 4c, d).

We can hypothesize how the CADH9-GAIN module is oriented relative to the membrane using our structure supplemented with AlphaFold2 models (Supplementary Fig. 4e, f)^76^. A large loop in the GAIN domain which is basic and hydrophobic could transiently interact with the outer surface of the membrane. Surfaces on EGF2/3, EGF8, and HormR also have surface exposed hydrophobic patches facing the putative plasma membrane. With this orientation, the flat plane of the CADH9-GAIN module is angled about 30 ° from the plasma membrane and the cadherin repeats would project outward at that angle.

### The CADH9-GAIN module is stabilized by interdomain contacts

Three key interfaces stabilize the CADH9-GAIN module (Fig. 2a, Supplementary Fig. 5, Supplementary Table 1). In the CADH9/GAIN interface (Fig. 2b), conserved residues F2188 and Y2250 interact with L1163, L1165, and L1187. Additionally, N1184 and R1185 from CADH9 interact with several residues on GAIN. Additional residues are involved, and this interface buries ∼1100 Å^2^ from solvent. Hydrophobic contacts are also seen between EGF2 and EGF7, burying ∼470 Å^2^ from solvent (Fig. 2c). I1379 and Y1383 of EGF2 pack against W1972, W1973, V1977, and P1980 of EGF7. A robust hydrophobic core is observed between LamG1 and EGF6, burying ∼770 Å^2^ from solvent (Fig. 2d). Residues V1581, F1585, Y1588, and V1589 from LamG1 pack against F1918, V1923, P1930, and Y1947 from EGF6. Also, D1578 interacts with the hydroxyl of Y1947.

**Fig. 2.**
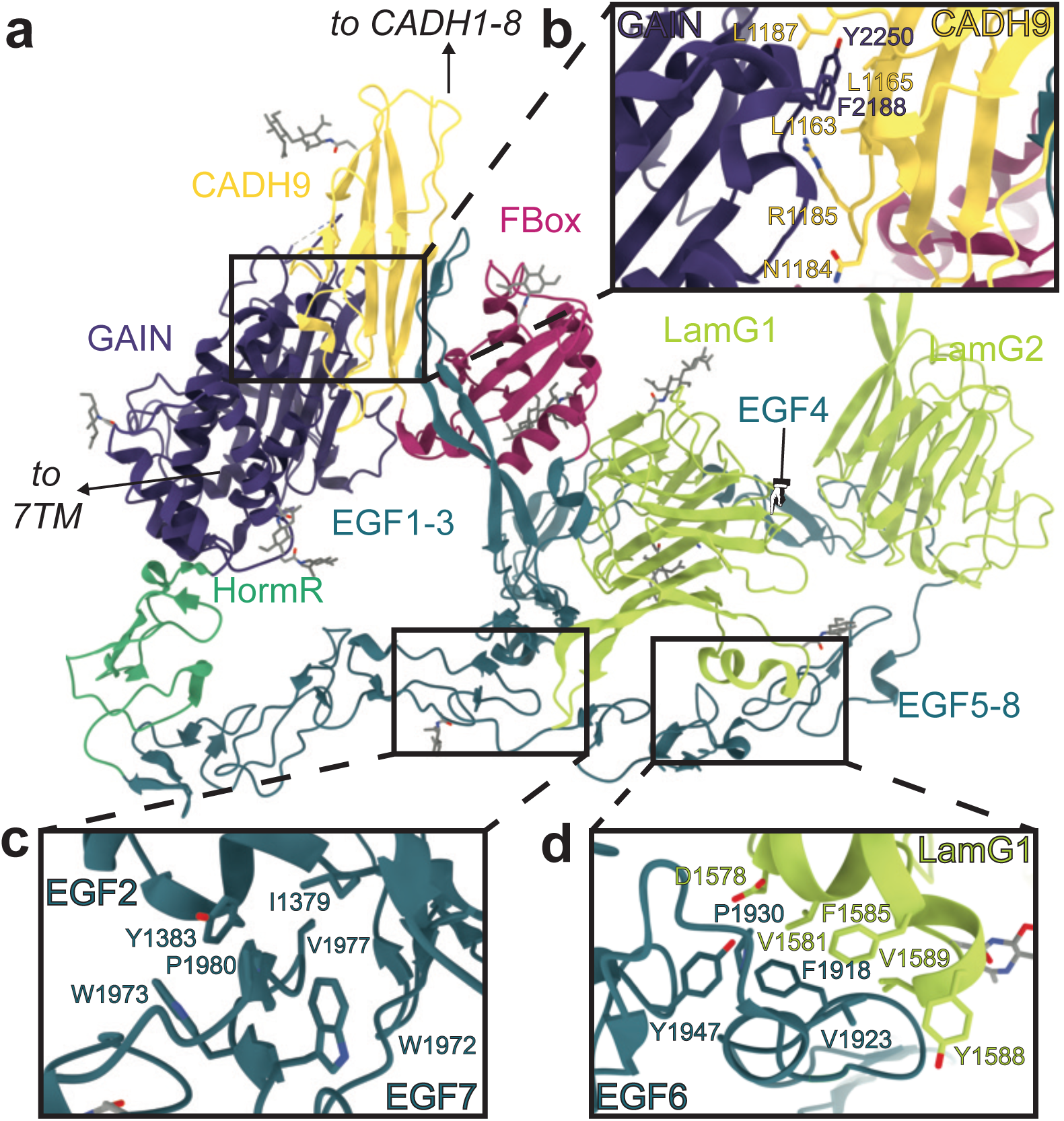
Structural details of interdomain interfaces stabilizing the CELSR1 ECR compact bundle. **a** Cartoon representation of CELSR1 ECR structure with key interfaces enclosed in black rectangles. **b** Cartoon model showing the interface between CADH9 and GAIN domains. Important residues are shown as sticks. **c** Cartoon model showing the interface between EGF2 and EGF7 domains with important residues shown as sticks. **d** Cartoon model showing the EGF6/LamG1 interface with important residues shown as sticks.

#### CELSR1 forms a Ca^2+^-dependent dimer with an extended conformation

We did not observe the CADH1-8 domains in our cryo-EM reconstruction even though these domains were about 40 % of the mass of our CELSR1 construct. CADH repeats have flexible interdomain linkers that rigidify upon Ca^2+^ binding, forming extended species that multimerize^46,47^. In order to resolve the CADH repeats, we biophysically characterized the CADH1-GAIN construct in the absence and presence of Ca^2+^ using inline size exclusion chromatography coupled to multi-angle light scattering and small angle X-ray scattering (SEC-MALS-SAXS) (Fig. 3, Supplementary Data 1, 2). We observed a leftward shift in the elution volume in the presence of 1 mM CaCl_2_ (Fig. 3a) and an increase in observed molecular mass by MALS, suggesting that the protein forms a dimer in solution. In the 1 mM CaCl_2_ condition, the main peak is likely a mixture of a dimer and a monomer due to its MALS molecular weight of 420 kD, which corresponds to roughly 60% dimer (∼ 520 kD) and 40 % monomer (∼ 260 kD) (Fig. 3a).

**Fig. 3.**
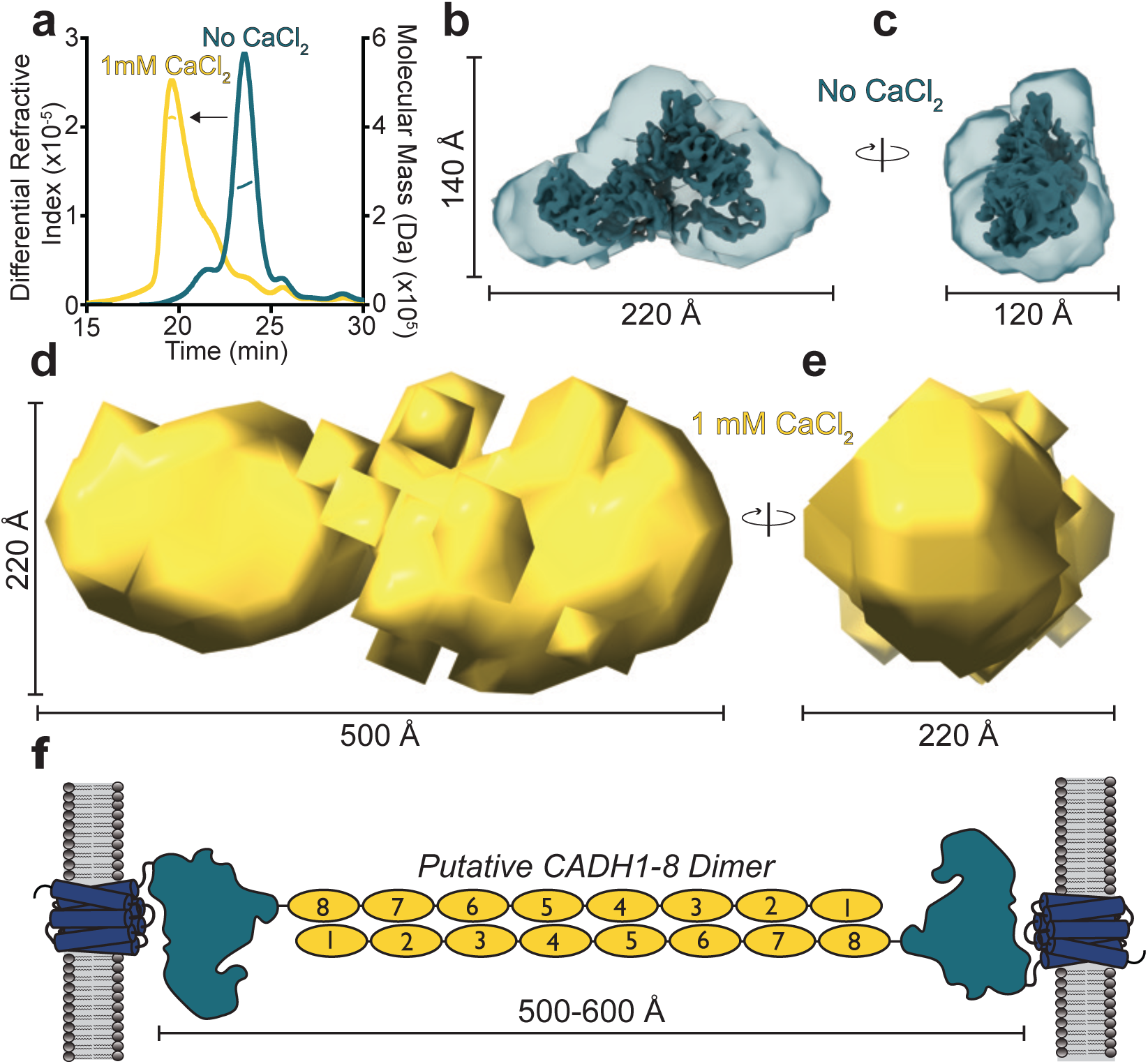
SEC-MALS-SAXS analysis of CELSR1 CADH1-GAIN. **a** Overlay of SEC-MALS traces in the absence of CaCl_2_ (blue curve) and in the presence of 1 mM CaCl_2_ (yellow curve). A leftward shift of the elution volume is seen with 1 mM CaCl_2_, suggesting the presence of a Ca^2+^-dependent dimer. Differential refractive index is plotted on the left and the calculated molecular mass is plotted on the right axis. The traces represent the differential refractive index whereas the horizontal lines in the peaks show the calculated molecular masses. **b** An electron density map generated using DENSS based on the No CaCl_2_ SAXS curve agrees well with the cryo-EM coulomb potential map, suggesting that the CADH9-GAIN CMM exists in solution. **c** An orthogonal view shows that the overall shape of the density is flat in one dimension relative to the other two. **d** A density map generated using DENSS based on the 1 mM CaCl_2_ SAXS curve shows a large, extended structure much longer in one dimension. **e** Orthogonal view of **d**. **f** A cartoon diagram with a hypothesis for how the CADH repeat region is arranged within the extended density observed in **d,e**. The model can extend from 500-600 Å depending on the curvature of the cadherin repeats. The density in **d,e** has a maximum of 500 Å in its longest dimension whereas the 1 mM CaCl_2_ dataset has a pairwise distance distribution function with D_max_ = 675 Å.

Using evolving factor analysis (EFA) deconvolution techniques, we separated individual scattering components from the SEC-SAXS curves and derived electron density reconstructions for these species (Fig. 3b-e and Supplementary Data 1, 2). We separated out the scattering for the monomer species (Supplementary Data 1, 2) and found that a density map for the no CaCl_2_ condition corresponds well in shape with the cryo-EM coulomb potential map with similar dimensions at ∼ 220 x 140 x 120 Å (Fig. 3b, c), suggesting that the CADH9-GAIN CMM is present in solution. We used EFA to separate out the scattering from the dimer component in the 1 mM CaCl_2_ condition (Supplementary Data 1, 2). We then generated an electron density map and found that the dimer component is extended at ∼ 500 x 220 x 220 Å (Fig. 3d, e) and we propose that this extended architecture corresponds to CADH1-8 in an antiparallel configuration (Fig. 3f), similar to that reported from atomic force microscopy^58^. A model of the entire CELSR ECR in this antiparallel configuration (Fig. 3f) could range from roughly 500 – 600 Å in maximum dimensions depending on the curvature of the CADH repeats. This is in overall agreement with the 1 mM CaCl_2_ dataset as it has a pairwise distance distribution function with D_max_=675 Å and a small peak ∼ 520 Å (Supplementary Data 1, 2).

#### MD simulations support a hinge region between CADH5-6

Using the deposited crystal structure of human CELSR1 CADH4-7, we investigated the dynamics of the proposed hinge region within the cadherin repeat region of CELSR^58,59^. In CELSR1, K533 and T564 between CADH5 and CADH6 are non-canonical. These positions are typically acidic calcium coordinating residues. These substitutions prevent the coordination of two out of three Ca^2+^ ions usually found between cadherin repeats. We performed three simulations (Fig. 4a), one with the wild-type CADH4-7 (WT), another where all calcium ions in the CADH5/CADH6 hinge were deleted (ΔCa^2+^), and a third where the non-canonical calcium coordinating residues were changed to the canonical ones, K533E/T564D, and Ca^2+^ ions were added manually (Canonical). We analyzed bending angles between CADH5 and CADH6 throughout each simulation and found that while the Canonical simulation remained close to 180° for the 100 ns simulation, both the WT and ΔCa^2+^ simulations diverged from this and displayed bending after 50 ns. We performed principal component (PC) analysis on these simulations and performed unbiased hierarchical clustering in the PC space (Fig. 4b). After obtaining three clusters (Cluster A-C), we examined frames of the simulation near the centroid of each cluster to understand the conformational dynamics (Fig. 4c). In Cluster A, the molecule is bent to ∼ 100° about the CADH5/CADH6 linker region. Cluster B is represented by a ∼ 135° bend in the CADH5/6 linker. Cluster C corresponds to an approximately linear configuration (180°). We quantified the percent of time that each variant spent in each cluster (Fig. 4d) and found that the WT and ΔCa^2+^ simulations can flex about the CADH5/6 hinge and sample both extended and bent conformations. However, these two simulations spend the majority of their time in bent conformations (Clusters A and B). In contrast, the Canonical simulation is largely limited to the extended conformation (Cluster C).

**Fig. 4.**
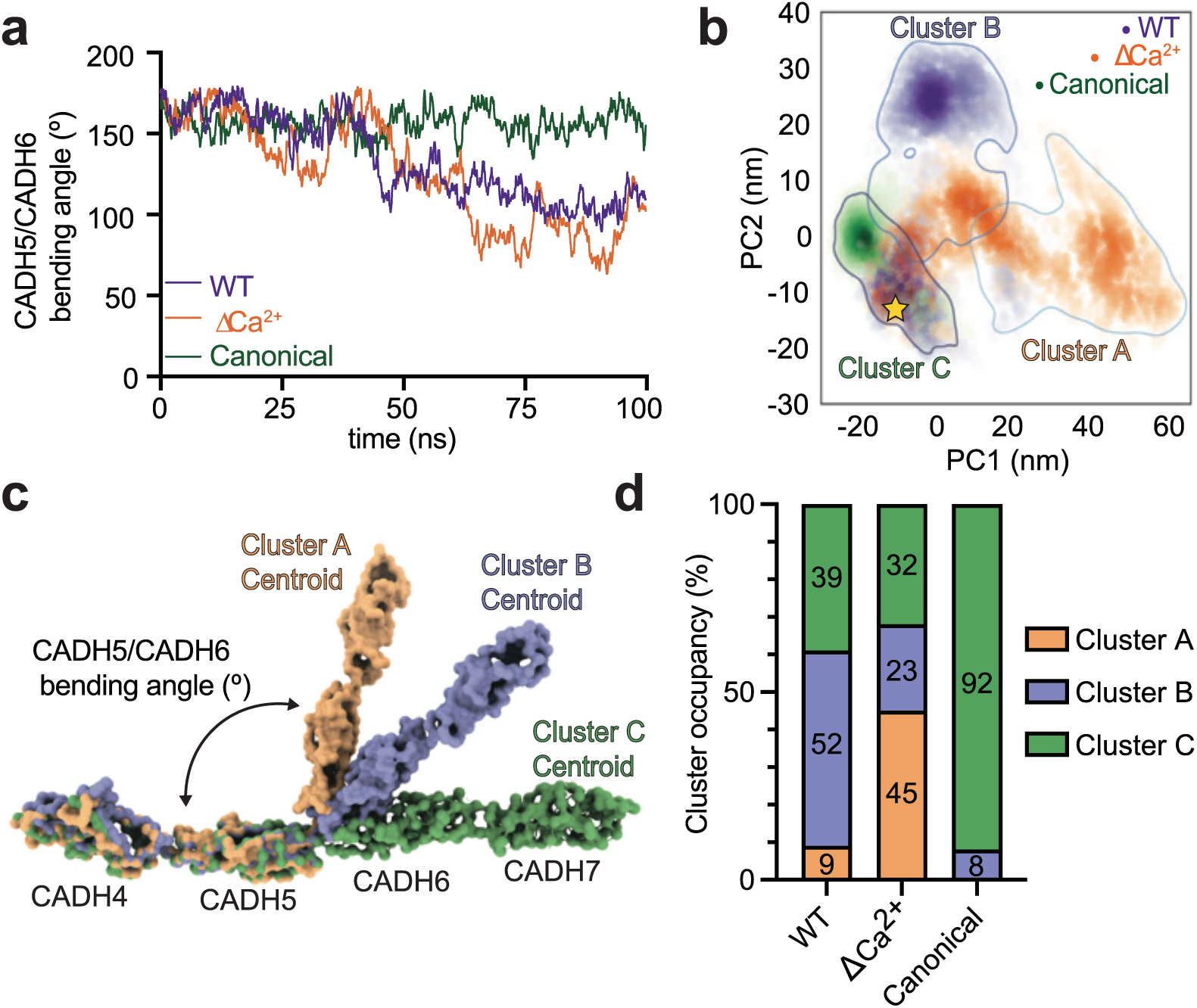
Molecular dynamics simulations on hCELSR1 CADH4-7 support that a flexible hinge exists between CADH5 and CADH6. **a** Plot of CADH5/CADH6 bending angle over time for each simulation (WT - purple, ΔCa2+ - orange, Canonical - green). **b** Kernel density estimates of each simulation plotted onto PC space. A yellow star shows the start point of each simulation. Clustering of conformations in essential principal component space reveals three clusters, labeled as clusters A-C. **d** Each shown PDB model is a frame of the simulation that is closest to the centroid of the designated cluster. **d** The percentage of simulation time that each simulation spends in each cluster.

#### CADH1-8 are essential for the adhesive functions of CELSR1

To address which portions of CELSR1 contribute to its adhesive and signaling functions, we split the protein into two modules according to our structural information. We deleted the CADH1-8 module (ΔCADH1-8) as well as the CMM resolved in our structure (ΔCADH9-GAIN) (Fig. 5a). We tested the expression of our constructs using fluorescence microscopy (Fig. 5b-d) and western blotting (Supplementary Fig. 6a-c) and used their relative cell-surface expression levels to normalize further experiments. We then profiled the adhesive function of our constructs in cell aggregation assays as well as a cell-cell junction enrichment assay. In the cell aggregation assay, each full-length protein is expressed on two different HEK293T cell populations and a fluorescent reporter is co-expressed. The cells are then mixed, and cell-cell aggregation is imaged and quantified^77^. WT CELSR1 was able to efficiently induce aggregation (Fig. 5e-g), the ΔCADH1-8 construct was defective in aggregation, and the ΔCADH9-GAIN construct was able to form aggregates but not as efficiently as WT. In the cell-cell junction enrichment assay, full length protein is expressed using HEK293T cells observed in groups, and the localization of CELSR1 is compared to ZO-1, a cell-cell junction marker^78,79^. Similar to the results for the cell aggregation assay, in the cell-cell junction assay, WT CELSR1 efficiently localized to the cell-cell junction (Fig. 5h-j) whereas the ΔCADH1-8 construct was found distributed all over the cell surface and not only at the cell-cell junction. The ΔCADH9-GAIN construct also localized to the cell-cell junction. We thus report the CADH1-8 module as sufficient for cell-cell adhesion. Neither deletion construct affected CELSR1 signaling activity (Supplementary Fig. 6d-e).

**Fig. 5.**
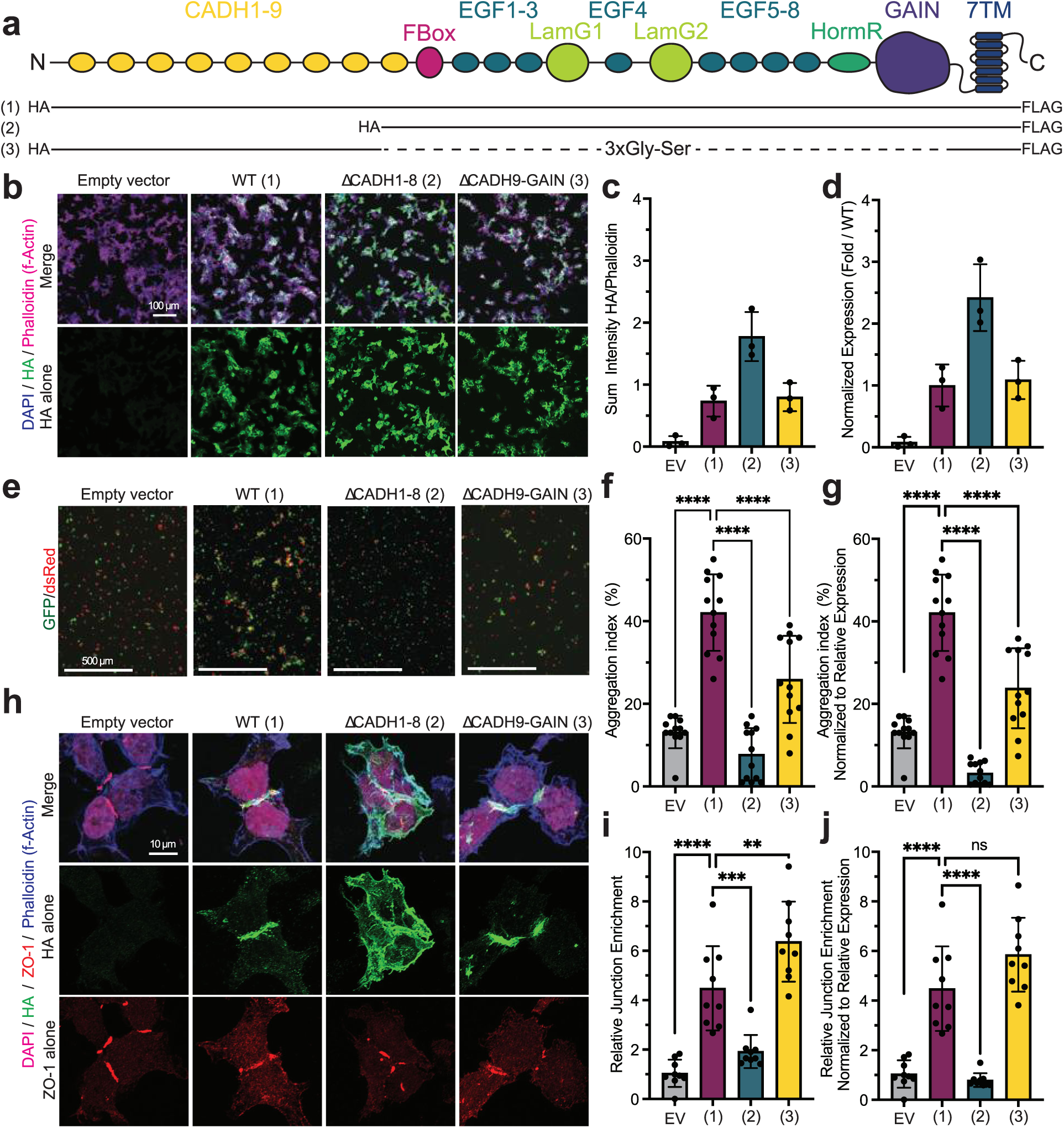
Functional analysis of the adhesive properties of the CELSR1 ECR. **a** Domain boundaries of constructs used in this study. Black solid lines show domain boundaries with dotted lines indicating regions that are connected with linkers. **b** Experimental constructs were tested for cell-surface expression using non-permeabilized cells imaged using the N-terminal HA tag (green). DAPI marks the nucleus (blue), and phalloidin marks filamentous actin (pink). **c** The ratio of HA/Phalloidin intensity is used to quantify relative cell-surface expression of each truncation construct. **d** The relative cell surface expression of each construct is presented as a fold change over wild type. **e** Representative cell-cell aggregation assay images. Two cell populations were transfected with either GFP or dsRed and the noted constructs, and cells were imaged to assess aggregation. **f** Quantification of cell aggregation images as aggregation index. N=3 independent biological experiments each done in quadruplicate; each data point is shown. The average is plotted as a bar graph with standard deviation shown for the error bars. **g** Quantification of aggregation index normalized to the relative cell surface expression of each construct. **h** Representative cell-cell junction enrichment assay images. Experimental constructs were imaged using the N-terminal HA tag (green). ZO-1 is used as a marker for the cell-cell junction (red), DAPI for the nucleus (pink), and phalloidin for filamentous actin (blue). **i** Quantification of images from **h** as relative junction enrichment. N=3 independent biological experiments each done in triplicate; graph as in **f**. **j** Quantification from **i** normalized to relative cell surface expression of each construct. one-way ANOVA with Tukey’s correction for multiple comparisons was performed to assess statistical significance between CELSR constructs. *** corresponds to p = 0.0001, **** corresponds to p < 0.0001. Each construct was compared to each other construct for the statistical testing, but only certain comparisons are shown for clarity.

We also found that WT CELSR1-mediated aggregation was abolished in the presence of EGTA, a selective Ca^2+^ chelator (Supplementary Fig. 6f-h) whereas aggregation mediated by TEN2/ADGRL3 was not affected by EGTA, serving as a control^61^. Finally, we found that CELSR1 and CELSR2 can form heterophilic aggregates when two cell populations expressing these proteins are mixed and this effect also is disrupted by EGTA supplementation (Supplementary Fig. 6f-h).

#### Mutations in the CADH9/GAIN interface enhance cellular adhesion

In order to understand the role of the CADH9-GAIN CMM in modulating CELSR function, we designed point mutations in the interface between CADH9 and GAIN from our cyro-EM structure. We designed three mutants targeting side chains that contribute to the interface: L1163A/F2188A (LF), L1163A/Y2250A (LY), and N1184A/R1185A/Y2250A (NRY). These mutants were tested for relative expression using fluorescence microscopy (Fig. 6 a-c) and their relative expression was used to normalize the cell aggregation data. We then performed cell aggregation assays where either empty vector or CELSR1 constructs were co-transfected with fluorescent proteins (Fig. 6d-f). We found that all three mutant constructs induced significantly higher aggregation in our assay than WT CELSR1, suggesting that if the CADH9-GAIN CMM is disrupted, the adhesive potential of CELSR1 is increased. We tested these point mutants in our signaling assay and found that none of our mutants had altered signaling activity compared to WT CELSR1 (Supplementary Fig. 7).

**Fig. 6.**
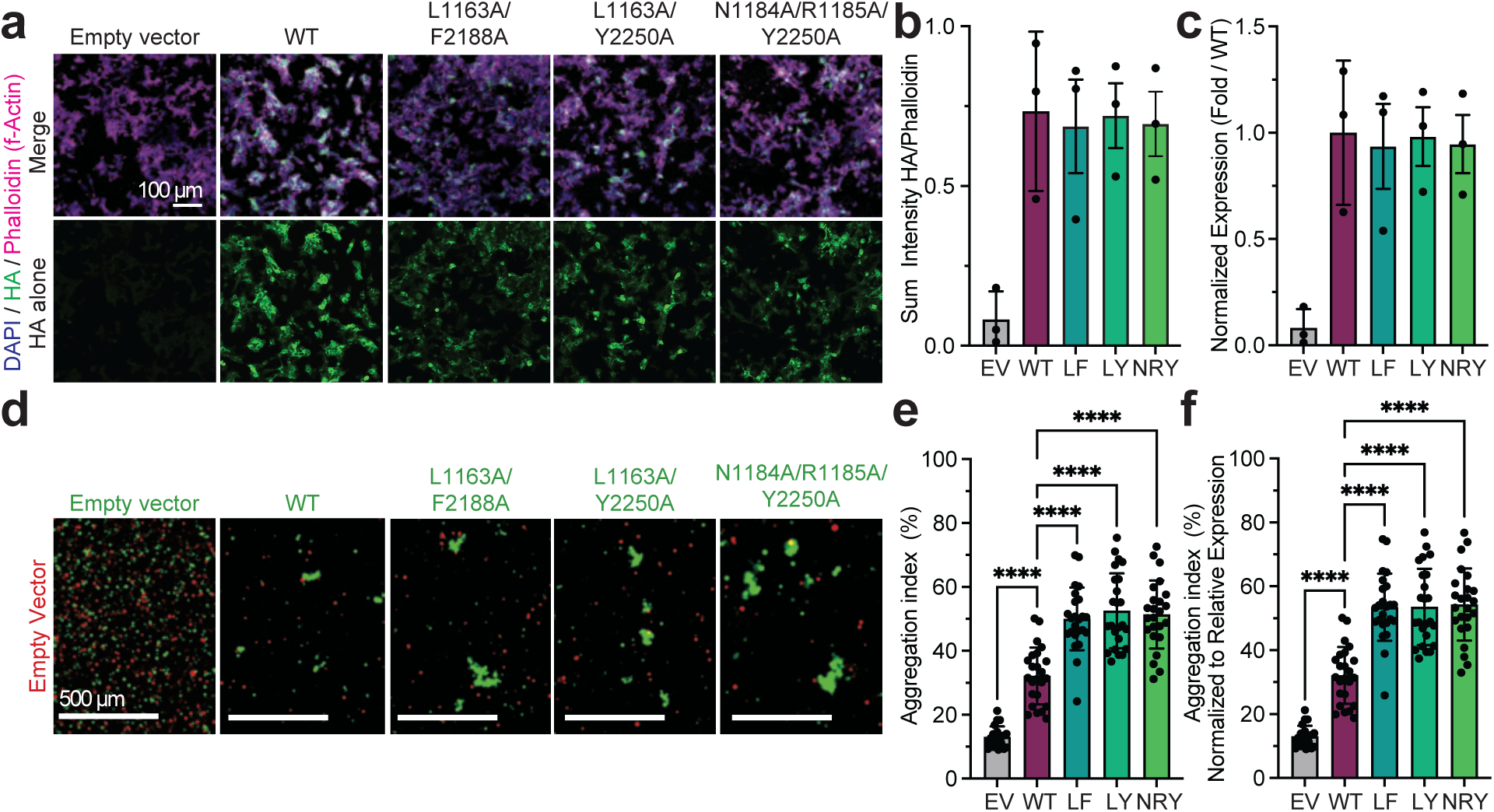
Point mutations in the CADH9/GAIN interface enhance CELSR1-mediated cell aggregation. **a** Experimental constructs were tested for cell-surface expression using non-permeabilized cells imaged using the N-terminal HA tag (green). DAPI marks the nucleus (blue), and phalloidin marks filamentous actin (pink). **b** The ratio of HA/Phalloidin intensity is used to quantify cell-surface expression of each construct. **c** The relative cell surface expression of each construct is presented as a fold change over wild type. **d** Representative cell-cell aggregation assay images. Two cell populations were transfected with GFP and the noted constructs, or dsRed with Empty Vector, and cells were imaged to assess aggregation. **e** Quantification of cell aggregation images as aggregation index. N=3 independent biological experiments each done in quadruplicate; each data point is shown. The average is plotted as a bar graph with standard deviation shown for the error bars. **f** Quantification of aggregation index normalized to the relative cell surface expression of each construct. one-way ANOVA analysis with Tukey’s correction for multiple comparisons was performed to assess statistical significance between CELSR constructs. *** corresponds to p = 0.0001, **** corresponds to p < 0.0001. Each construct was compared to each other construct for the statistical testing, but only certain comparisons are shown for clarity.

## Discussion

CELSRs are highly conserved adhesion GPCRs critical for animal development^4–6^. Their large extracellular regions have remained enigmatic; their role in cellular adhesion, signaling, and other functions remain unclear, and molecular-level detail has been sparsely available. In this work, we determined the cryo-EM structure of the CELSR1 ECR, revealing a CMM containing CADH9-GAIN. We also used functional assays to define two modules of the protein; CADH1-8 is required for cellular adhesion whereas CADH9-GAIN regulates cellular adhesion.

Our work adds a key structure to the landscape of the adhesion GPCRs by defining the three-dimensional structure of the CELSR/ADGRC ECR. The CADH9-GAIN region of CELSR1 contains 14 domains arranged in a CMM mediated by conserved hydrophobic contacts between the N-terminal CADH9 domain and C-terminal GAIN domain. As such, the CADH9-GAIN CMM is likely present across CELSR/Flamingo orthologs. Our data provide molecular detail to a previous report showcasing low resolution atomic force microscopy images of the full CELSR2 extracellular region^58^. The CADH9-GAIN CMM is large, with a solvent-accessible surface area of 71,000 Å^2^ and presents several putative ligand binding sites to the extracellular milieu, including the putative ‘hypervariable’ surface of the LamG2 domain^72^. Others have suggested a role for this CADH9-GAIN CMM in the interaction of Fmi with Frizzled^70^, but the molecular details of this interaction, as well as the interaction of the known PCP ligands Frizzled and Van Gogh with CELSR1 in cis^13,80^, remain unclear^70^. The CADH9-GAIN CMM of CELSR3 is the proposed binding site for β-amyloid oligomers^81^. Additionally, CELSR2 has been shown to dimerize in a parallel configuration via its globular CADH9-GAIN CMM and this binding site remains to be determined^58^.

Evidence collected by us and others is consistent with a model of the CADH1-8 mediated antiparallel dimer driving the adhesive activity of CELSRs^13,14,21,70,82^. In the presence of Ca^2+^, we determined an extended species ∼ 500 Å long from our SAXS data which is consistent with an extended antiparallel dimer. This agrees with another report which used atomic force microscopy to determine low resolution images of CELSR ECR constructs^58^. In this model, each protomer is twisted around the other, where CADH1 interacts with CADH8 on the opposing monomer, CADH2 with CADH7, and so forth. This dimerization mode is similar to known protocadherin structures, where the cadherin repeats are arranged in an antiparallel fashion, and they twist around each other^52–57^. We propose that this trans antiparallel dimer mediates the adhesive function of CELSR, as our deletion of the CADH1-8 module abrogates cell aggregation activity as well as the cell-cell junction localization of CELSR1 in cells. The use of EGTA disrupted WT CELSR1-mediated aggregation, also consistent with a Ca^2+^-dependent antiparallel dimer of CADH repeats. These data are consistent with what others have found using the cell aggregation assay^13,14,21^, including a group which recently found that CADH1-8 of the CELSR homolog Fmi is necessary and sufficient for cell adhesion^70^. This tight multi-CADH dimer thus may be relevant to the role that CELSRs play in many processes which require the maintenance or coordinated change of cell-cell contacts in response to changes in force or pressure^6,9,11,17,18^. Future work will reveal the detailed molecular architecture of the CELSR antiparallel dimer, explaining the mechanism of how the known CELSR mutations Crash and Spin Cycle result in neuromuscular dysfunction^9^.

We propose that the CELSR1 CMM is the crucial structural feature of CELSR-subfamily aGPCRs (Fig. 7). Our data suggest that the CMM regulates the orientation and exposure of the adhesive CADH1-8 region in the extracellular space. For other adhesion molecule complexes, the rigidity of the ECR, the relative orientation between domains, is important for productive adhesion^83^. Point mutations in the CMM, which likely disrupt the CADH9/GAIN interface and could favor an extended CMM conformation, enhance CELSR1 mediated cell aggregation. This is consistent with a model proposed for Fmi, where binding of Frizzled to Fmi in cis results in a conformational change that enhances the CADH1-8 dimer in trans^70^. The ability of the CELSR CMM to transition between the compact conformation we observe in our cryo-EM structure, and a putative extended conformation (driven by our point mutations or by a binding partner), may explain how the symmetric CADH1-8 dimer can be regulated in an asymmetric fashion in order to establish PCP. This is similar in principle to the closed/compact and open/extended conformations observed for the ADGRG6 ECR, which are regulated by alternative splicing^84^. We speculate based on this work that aGPCRs, in general, may have N-terminal regions of their ECRs as adhesive regions and that the adhesive signal is communicated to a CMM, which can regulate this adhesive potential. Indeed, several aGPCRs employ their N-terminal domains to bind a protein ligand: ADGRL3 with TEN2 and FLRT^61,85–87^, ADGRG1 with transglutaminase 2^88^, and ADGRG6 with Collagen IV^89^. We add to this list CELSR1 which uses its N-terminal CADH repeats to mediate homophilic adhesion in trans. In each case, the N-terminus extends farthest away from the aGPCR-expressing cell, so it is poised to interact with ligands in the extracellular space. The adhesive ‘signal’ can then be transferred to a CMM, the simplest version of which is the GAIN domain alone. Changes in CMM conformation could also dictate which binding partners could be recruited in cis, or binding partners in cis could change CMM conformation. It is also possible that this N-terminal adhesive signal operates through the CMM to regulate downstream receptor function, although for CELSRs it remains to be seen how signaling activity may relate to PCP function^90^.

**Fig. 7.**
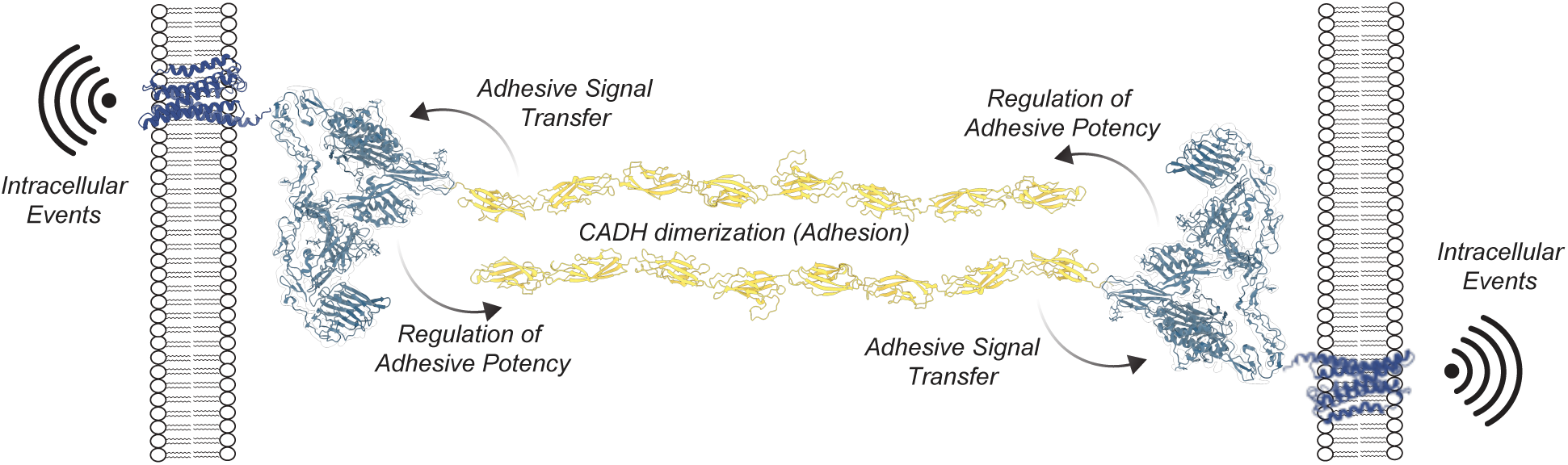
Integrative model for ECR-mediated functions of CELSR1. The dimerization of cadherin repeat region, CADH1-8, leads to cell aggregation and localization to the cell-cell junction. This adhesion can be regulated by the CADH9-GAIN CMM. The receptor can induce intracellular events in response to these adhesive events in the extracellular space.

Additionally, other aGPCRs have CMMs. ADGRG6 has 5 domains in a CMM mediated by its N-terminal CUB domain^75^. ADGRG1 has an N-terminal PLL domain which is found tightly packed against its GAIN domain, generating a small, prototypic CMM^91^. In both ADGRG6 and ADGRG1, splice variants can disrupt these CMMs in order to regulate ligand binding and receptor function^75,91,92^. Overall, the intricate, modular architectures of aGPCRs likely evolved to withstand and react to strict spatial and force requirements in order to mediate intracellular events^7,93–98^. More experimental structures of aGPCRs with larger extracellular regions, such as ADGRV1, as well as comprehensive mechanistic studies including structural biology, adhesion, and functional readouts, will determine whether the CMM is a general structural feature of adhesion GPCRs.

## Materials and Methods

### Cloning and construct design

All amino acid numbering in this text is based on *M. musculus* CELSR1, UniProt entry O35161. Bioinformatic tools including PROMALS3D^99^, SMART^100^, PSIPRED^101^ and CDD^102^ were used to identify domain boundaries and design CELSR1 constructs. The full length CELSR1 and CELSR2 constructs in the pEB vector with N-terminal HA and C-terminal FLAG tags were previously described^103^. A region containing the CELSR1 CADH1 through GAIN domains (residues 255-2477) was amplified with these primers: F: 5’-CTGCCTTTGCGGCGAGCACCTCCCCACAGTTCCCCCTGCC-3’ and R: 5’-GTGGT GATGGTGATGATGATGATGCTCCCGTCTGCTTATGTC-3’ and inserted into the pAcGP67a vector with a C-terminal 8 x histidine tag using Gibson assembly^104^. The ΔCADH9-GAIN construct was generated with construct was generated using Gibson assembly by digesting the pEB backbone with EcoRV-HF and BamHI-HF (New England Biolabs) and inserting 2 Gibson fragments amplified using the following primers: 5’-cggtatcgataagcttgatatcgaattcct-3’ and 5’- actgccactaccgctgccCTGGTCCAGGAGACGGATGT-3’; 5’- cagggcagcggtagtggcagtGACATTTCCAGACGTGAGCACG-3’ and 5’-gccgctctagaactagtggatcc-3’. The ΔCADH1-8 construct was generated using Gibson assembly by digesting the pEB backbone with BamHI-HF (New England Biolabs) and inserting 3 Gibson fragments amplified using the following primers: 5’-aattcctgcagcccgggggatc-3’ and 5’-agcatagtcaggtacatcataggggtaagcaactgcag-3’, 5’- gatgtacctgactatgctTTGCCTGACTTCCAGATCCTTTTCAACAACTATG-3’ and 5’- gCGTCATTCATGTTCAGGGTGGCaatgttg-3’, 5’-GCCACCCTGAACATGAATGACGc and 5’-gccgctctagaactagtggatccTCAtttatcg-3’. The L1163/F2188 mutant was generated using pEB digested with BamHI and a four-part Gibson assembly using fragments generated with the following primers: 5’- aattcctgcagcccgggggatc-3’ and 5’- cgggatccagcagcagcaagctcgcctcgttgccttgc-3’, 5’- gcaaggcaacgaggcgagcttgctgctgctggatcc-3’ and 5’-gctgctaggtcagcgccctgctggcggc-3’, 5’- gccgccagcagggcgctgacctagcagcc-3’ and 5’-gccgctctagaactagtggatccTCAtttatcg-3’. The L1163/Y2250 mutant was generated using pEB digested with BamHI and a four-part Gibson assembly using fragments generated with the following primers: 5’-aattcctgcagcccgggggatc-3’ and 5’- cgggatccagcagcagcaagctcgcctcgttgccttgc-3’, 5’-gcaaggcaacgaggcgagcttgctgctgctggatcc-3’ and 5’- ggtgacgatgacgaagggcctcagagcggtcctcttcac-3’, 5’-gtgaagaggaccgctctgaggcccttcgtcatcgtc-3’ and 5’- gccgctctagaactagtggatccTCAtttatcg-3’. The N1184/R1185/Y2250 mutant was generated using pEB digested with BamHI and a four-part Gibson assembly using fragments generated with the following primers: 5’-aattcctgcagcccgggggatc-3’ and 5’- CATGAGCGCCTCCAGTGGagctgcGTTGTCCAGATCCCGGCTGA-3’, 5’- CAGCCGGGATCTGGACAACgcagctCCACTGGAGGCGCTCATG-3’ and 5’- ggtgacgatgacgaagggcctcagagcggtcctcttcac-3’, 5’-gtgaagaggaccgctctgaggcccttcgtcatcgtc-3’ and 5’- gccgctctagaactagtggatccTCAtttatcg-3’.

#### Cell Culture

Sf9 cells (Thermo Fisher, 12659017) were cultured in SF900-III medium with 10% (v/v) FBS (Sigma-Aldrich, F0926) at 27 °C and transfected with plasmids and commercial baculovirus DNA (Expression Systems, 91-002) using Cellfectin II (Thermo Fisher, 10362100). Following transfection, initial baculoviral stocks were harvested and Sf9 cells were used produce high-titer recombinant baculovirus. As previously described^61^, High Five cells (*Trichoplusia ni*, female, ovarian ThermoFisher, B85502) were used for production of recombinant proteins. High Five cells were cultured using Insect-Xpress medium (Lonza, 04351Q) with 10 μg/mL gentamicin at 27 °C. High Five cells (Thermo Fisher, B85502) were infected with baculovirus at 2.0 x 10^6^ cells/mL and incubated at 27 °C with 120 rpm shaking for 72 h.

HEK293T mammalian cells (ATCC, CRL-3216) were used for cell aggregation assays similar to previously described^77^ and were cultured in Dulbecco’s modified Eagle’s medium (DMEM; Gibco, 11965092) supplemented with 10 % FBS (Sigma-Aldrich, F0926) at 37 °C in 5 % CO_2_. HEK293T cells (ATCC #CRL-11268) were used for junctional enrichment studies. Cells for junctional enrichment were maintained in DMEM (Gibco Cat# 11995065) plus 10 % FBS (Gibco Cat# 16000044) and 1X Penicillin-Streptomycin (Corning Cat# MT30002Cl) at 37 °C and 5 % CO_2_ for a maximum of 20 passage numbers.^106,107^

#### Protein expression and purification

As previously described^61^, media was harvested from High Five cell culture 72 hours after baculovirus infection. The media was centrifuged at room temperature at 900 x g for 15 min. The supernatant was harvested, transferred to a beaker with stirring at room temperature, and the following were added: 50 mM Tris pH 8.0, 5 mM CaCl_2_ and 1 mM NiCl_2_ (final concentrations listed). After 30 min, the solution was centrifuged for 30 min at 8000 x g. The clarified supernatant was then incubated with nickel-nitriloacetic (Ni-NTA) resin (QIAGEN 30250) with stirring at room temperature for at least 3 hours. A Büchner funnel was used to collect resin and wash using a buffer composed of 10 mM Tris pH 8.5, 150 mM NaCl (TBS) with 20 mM imidazole, and then the washed resin was transferred to a poly-prep chromatography column (Bio-Rad). The protein was eluted using TBS buffer plus 200 mM imidazole.

Fractions containing desired protein were pooled and concentrated using a 100 kDa centrifugal concentrator (Amicon UFC810024) and loaded on gel filtration chromatography using a Superose 6 Increase 10/300 column (GE Healthcare) using the following buffer: 10 mM Tris pH 8.5, 150mM NaCl, with the supplementation of 1 mM CaCl_2_ as necessary.

#### SAXS data collection and processing

See Table 2 for data collection and processing statistics, as well as Supplementary Data files 1 and 2. SAXS was performed at BioCAT (Beamline 18ID at the Advanced Photon Source, Chicago) with in- line size exclusion chromatography (SEC) to separate sample from aggregates and other contaminants thus ensuring optimal sample quality and multiangle light scattering (MALS), dynamic light scattering (DLS) and refractive index measurement (RI) for additional biophysical characterization (SEC-MALS-SAXS). The samples were loaded on a Superose 6 Increase column (Cytiva) run by a 1260 Infinity II HPLC (Agilent Technologies) at 0.6 mL/min. The flow passed through (in order) the Agilent UV detector, a MALS detector and a DLS detector (DAWN Helios II, Wyatt Technologies), and an RI detector (Optilab T-rEX, Wyatt). The flow then went through the SAXS flow cell. The flow cell consists of a 1.0 mm ID quartz capillary with ∼20 μm walls. A coflowing buffer sheath is used to separate sample from the capillary walls, helping prevent radiation damage^105^. Scattering intensity was recorded using an Eiger2 XE 9M (Dectris) detector which was placed 3.6 m from the sample, giving access to a q-range of 0.003 Å^-1^ to 0.42 Å^-1^. 0.6 s (no CaCl_2_) or 0.4 s (1 mM CaCl_2_) exposures were acquired every 1 s during elution and data were reduced using BioXTAS RAW 2.1.4^106^. Buffer blanks were created by averaging regions flanking the elution peak and the entire series was buffer subtracted. The buffer subtracted series was analyzed using evolving factor analysis within RAW and separated into individual scattering components and the separated I(q) vs q curves were used for subsequent analyses. The GNOM and DENSS packages within RAW were used to calculate IFTs and electron density maps for each scattering component. Molecular weights and hydrodynamic radii were calculated from the MALS and DLS data respectively using the ASTRA 7 software (Wyatt).

**Table 2.**
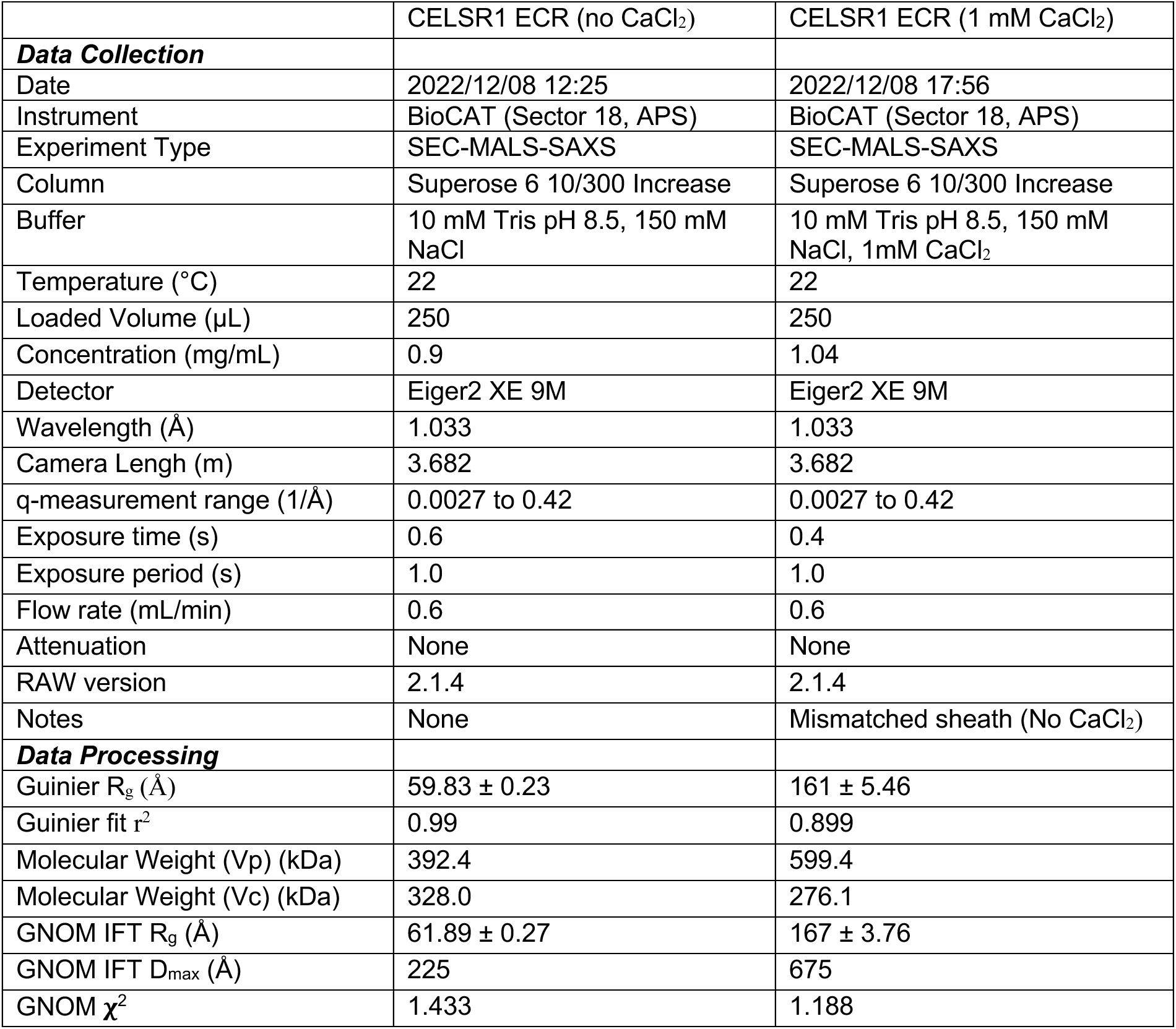
Small angle X-ray scattering data collection and data processing information.

|  | CELSR1 ECR (no CaCl<sub>2</sub>) | CELSR1 ECR (1 mM CaCl<sub>2</sub>) |
| --- | --- | --- |
| **Data Collection** |  |  |
| Date | 2022/12/08 12:25 | 2022/12/08 17:56 |
| Instrument | BioCAT (Sector 18, APS) | BioCAT (Sector 18, APS) |
| Experiment Type | SEC-MALS-SAXS | SEC-MALS-SAXS |
| Column | Superose 6 10/300 Increase | Superose 6 10/300 Increase |
| Buffer | 10 mM Tris pH 8.5, 150 mM NaCl | 10 mM Tris pH 8.5, 150 mM NaCl, 1mM CaCl<sub>2</sub> |
| Temperature (°C) | 22 | 22 |
| Loaded Volume (μL) | 250 | 250 |
| Concentration (mg/mL) | 0.9 | 1.04 |
| Detector | Eiger2 XE 9M | Eiger2 XE 9M |
| Wavelength (Å) | 1.033 | 1.033 |
| Camera Length (m) | 3.682 | 3.682 |
| q-measurement range (1/Å) | 0.0027 to 0.42 | 0.0027 to 0.42 |
| Exposure time (s) | 0.6 | 0.4 |
| Exposure period (s) | 1.0 | 1.0 |
| Flow rate (mL/min) | 0.6 | 0.6 |
| Attenuation | None | None |
| RAW version | 2.1.4 | 2.1.4 |
| Notes | None | Mismatched sheath (No CaCl<sub>2</sub>) |
| **Data Processing** |  |  |
| Guinier R<sub>g</sub> (Å) | 59.83 ± 0.23 | 161 ± 5.46 |
| Guinier fit r<sup>2</sup> | 0.99 | 0.899 |
| Molecular Weight (Vp) (kDa) | 392.4 | 599.4 |
| Molecular Weight (Vc) (kDa) | 328.0 | 276.1 |
| GNOM IFT R<sub>g</sub> (Å) | 61.89 ± 0.27 | 167 ± 3.76 |
| GNOM IFT D<sub>max</sub> (Å) | 225 | 675 |
| GNOM χ<sup>2</sup> | 1.433 | 1.188 |

#### Cryo-EM data collection

3.5 μL purified CELSR1 CADH1-GAIN (3.0 mg/mL) was applied on glow-discharged holey carbon grids (Quantifoil R1.2/1.3, 300 mesh), and vitrified using a Vitrobot Mark IV (FEI Company). The specimen was visualized at the National Cryo-EM Facility using a Titan Krios electron microscope (FEI) operating at 300 kV and equipped with a K3 direct electron detector (Gatan, Inc.) with the Latitude software (Gatan, Inc.). Images were recorded with a nominal magnification of ×81,000 in super-resolution counting mode, corresponding to a pixel size of 0.55 Å on the specimen level. To maximize data collection speed while keeping image aberrations minimal, image shift was used as imaging strategy using one data position per hole with six holes targeted in one template with one focus position. In total, 5826 images with defocus values in the range of −1.0 to −2.0 μm were recorded using a dose rate of 6.2 electrons/s/physical pixel. The total exposure time was set to 2.5 s resulting in an accumulated dose of about 50 electrons per Å^2^ and a total of 50 frames per movie stack. To improve resolution, a second dataset of 5550 micrographs was collected in exactly the same manner on the same microscope and the two particle stacks were combined at the stage of 3D classification.

#### Cryo-EM processing and model building

A visual representation of the data processing pipeline is available in Supplementary Fig. 2 and important values and statistics are available in Table 1. RELION 4.0 was used for the initial steps of data processing on each set of micrographs separately^107^. Stack images were subjected to beam-induced motion correction using MotionCor2 and binned to 1.1 Å/pixel^108^. The motion-corrected micrographs were then imported into CryoSPARC 4.1.2^109^ where all subsequent data processing occurred. CTF parameters for each micrograph were estimated using Patch CTF. 2,129,894 particles were picked using the Blob picker and subjected to reference-free two-dimensional classification to discard particles categorized in poorly defined classes. A refined set of 162,975 particles was used to train the Template picker which generated 1,794,239 particles. Again, reference-free two-dimensional classification was used to discard particles categorized in poorly defined classes and a final set of 79,985 particles were used in the Ab-Initio Reconstruction job with three classes which generated three volumes. The same three volumes were used in heterogeneous refinement with a larger particle stack, combining picks from the Blob and Template pickers (removing duplicates) to generate a higher quality volume. This volume had 235,891 particles associated with it and these particles were used to train the Topaz picker which picked 1,040,008 particles. Again, reference-free two-dimensional classification was used to discard particles categorized in poorly defined classes and a final set of 947,890 particles was merged with the picks from Autopicking and Template picking with duplicates removed. Three volumes were used in multiple rounds of heterogenous refinement with merged particles until the map quality no longer improved, and Non-Uniform Refinement was used to obtain the final coulomb potential map.^78,113^ Reported resolutions are based on the gold-standard Fourier shell correlation (FSC) using the 0.143 criterion^110^. Model building started from the AlphaFold2 model of CADH9-GAIN, which was docked into the EM density map using Phenix, refined using the real-space refinement module in the Phenix software suite^111^ and then manually checked and adjusted residue-by-residue in an iterative fashion along with further real-space refinement to fit the density using COOT^112^. The final model contains residues 1113-2470 and the final model statistics are provided in Table 1.

#### Structural analysis and sequence conservation analysis

Structural analysis and manual inspection of structures was performed using PyMOL version 2.4.1 (Schrodinger) and sequence analysis was done using PROMALS3D ^99^ and the ConSurf^113^ server. ChimeraX was used to analyze cryo-EM density maps and generate figures^114^.

#### Molecular dynamics simulations

The structure used for these simulations was previously solved by Tamilselvan and Sotomayor (PDB ID 7SZ8)^59^. Atoms missing from the crystal structure were added with Swiss PDB Viewer. Crystal structure waters and sodium ions were removed from the protein. Using GROMACS version 2022.1, CELSR1 CADH4-7 was placed in an 8216.1 nm^3^ rhombal dodecahedron with side length 13.9 nm and periodic boundary conditions, ensuring over 1 nm of solvent between the most distant atoms of 7SZ8 and the boundaries. The box was solvated with the TIP3P water model using gmx solvate and Na^+^ and Cl^−^ counterions were added using gmx genion. The protein structure was minimized with the leapfrog integrator in step sizes of 0.1 Å until the max force was smaller than 1000 kJ mol^-1^ nm^-1^.

The system was then subjected to a two-phase equilibration in the NVT and NPT ensembles with protein atoms under harmonic position restraints. A 100 ps NVT equilibration was carried out with step size 2 fs at 300 K with the Berendsen thermostat using a 0.1 ps coupling constant, and solvent velocities were assigned from the Boltzmann distribution. NPT equilibration was also carried out with step size 2 fs for 100 ps, and at 1 atm with Parrinello-Rahman pressure coupling (2.0 ps coupling constant) using velocities from NVT ensemble. Simulations on the unrestrained protein were run in the NPT ensemble for 1 μs and a 2 fs step size. Long-range electrostatics were computed with PME with a 1.6 Å Fourier spacing and 10 Å cutoff. All simulations were run with the AMBER99SB-ILDN force field. Different simulation conditions were separately solvated, minimized, and equilibrated as above.

### Simulated systems

| Condition | Simulation Length |
| --- | --- |
| WT | 100 ns |
| K533E/T564D<br>(Canonical) | 100 ns |
| $\Delta\text{Ca}^{2+}$ | 100 ns |

The WT simulation was prepared as described above. The K533/T564 simulation was designed by aligning calcium coordinating junctions (CADH 4-5 and CADH 6-7) to the CADH 5-6 junction. After alignment, K533E and T564D were changed to match the calcium coordination geometry of the canonical sites, and the two missing calcium ions were added into their respective positions to make a three-calcium interdomain junction between cadherins 5 and 6. The ΔCa^2+^ simulation was prepared by removing the calcium ion between CADH 5 and 6 from the WT protein.

### PCA and free energy landscape analysis

Trajectories from the simulations were extracted for the alpha carbons (Cαs) of each residue for each simulation. Principal components of the Cαs were calculated by concatenating all three simulation trajectories for each with gmx trjcat and diagonalizing the covariate matrix with gmx covar. Trajectories were projected onto essential gmx anaeig and were plotted and visualized with a homemade python script. The fractions of variance ! captured by PCs were checked to verify that the first two principal components captured most of the variance and verify the quality of the dimension reduction by computing

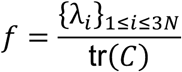

where {λ_i_}_1≤i≤3N’_ is the set of 3N eigenvalues for N α-carbons, and tr(C) is the trace of the covariate matrix. Hierarchical clustering of free energy landscape wells was performed by scipy.cluster in a homemade python script using the Ward clustering method to minimize variance in the l^2^ norm. Cluster centroids c_i_ were computed by averaging the PC1 and PC2 coordinates of frames within each cluster

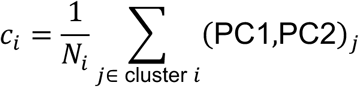

where N_i_ is the number of frames in the i^th^ cluster. We chose the frame from the simulations whose PC coordinates are closest by l^2^ norm and constructed the 3D structure as a linear combination of PC1 and PC2 using gmx anaeig.

#### Cell aggregation assay

As previously described^77^, HEK293T cells (ATCC, CRL-3216) were seeded using a 6-well plate in 2 mL of DMEM supplemented with 10% FBS and incubated overnight at 37 °C. When the cells reached ∼ 60 % confluency, they were co-transfected with 1 µg each of either pCMV5 + GFP or dsRed, CELSR1 + GFP or dsRed, CELSR2 + GFP or dsRed, ADGRL3 + GFP and TEN2 + dsRed and 5 μL LipoD293T (SL100668; SignaGen Laboratories) or Fugene 6 (Promega E2691). Two days after transfection, the media was aspirated and the cells were washed with 1xPBS, detached with 1xPBS containing 1 mM EGTA, and supplemented with 15 µL of 50 mg ml^−1^ DNAse IV (Sigma, D5025). The cells were resuspended by pipetting to a create single-cell suspensions, transferred to a microcentrifuge tube and an additional 15 µL of DNAse solution was added to each sample. Seventy µL of cells expressing the indicated constructs were mixed in a 1:1 ratio in one well of a non-coated 24-well plate containing 360 µL of DMEM supplemented with 10 % FBS, 20 mM CaCl_2_ and 10 mM MgCl_2_ or DMEM with 10 % FBS, 10 mM MgCl_2_, and 20 mM EGTA for 5 hours at 126 rpm at 37 °C with 5% CO2 and imaged using a Leica Fluorescence Microscope with a 5× objective. Aggregation index at time = 5 hrs was calculated using ImageJ 1.52e. A value for particle area of 2-3 cells was set as a threshold based on negative control values. The aggregation index was calculated by dividing the area of particles exceeding this threshold by the total area occupied by all particles in the individual fields.

#### Immunocytochemistry

Cover glass (#0, 12 mm, Carolina Biological Supply Company #633009) was placed into 24-well plates and coated for 2 hrs with 100 µL of 50 µg/mL poly-D-lysine (Gibco #A38904-01) in the 37 °C tissue culture incubator. Excess poly-D-lysine was removed, coverslips were washed 3x with sterile ddH_2_O, and dried for 30 mins. HEK293T cells were plated at 1.5 x 10^5^ cells/well in 0.5 mL complete DMEM. After 16-24 hrs, cells were transfected with indicated experimental plasmid via TransIT-2020 (Mirus MIR5400) with a total of 0.5 µg DNA amount/condition/well. After 48 hrs post-transfection, cells were washed briefly once with PBS, fixed with 4 % PFA (Electron Microscopy Science Cat# 15714) with 4% sucrose in PBS for 20 min at 4 °C, and washed 3 x 5 mins in PBS. For surface receptor labeling of HA tag, samples were then transferred directly into blocking buffer (4 % BSA (Sigma Cat# 10735086001) plus 3 % normal goat serum (Jackson Immunoresearch #005000121) in PBS) for 1 hr. Samples were then labeled with HA tag primary antibody (anti-HA mouse, Covance Cat# MMS101R; 1:2,000) diluted into blocking buffer for 2 hrs at room temperature. Samples were washed 3 x 5 mins with PBS, permeabilized with 0.2% Triton X-100 in PBS for 5 mins at room temperature and incubated in primary ZO-1 antibody (ZO-1 polyclonal rabbit antibody, Invitrogen #61-7300, 1:2,000) diluted into blocking buffer for 2 hrs at room temperature. Samples were subsequently washed 3 x 5 mins with PBS and incubated in fluorescently conjugated secondary antibodies (goat anti-mouse Alexa Fluor 488, ThermoFisher #A11001, 1:1,000; goat anti-rabbit Alexa Fluor 546, ThermoFisher #A11010, 1:1,000) diluted into blocking buffer together with Phalloidin Alexa 647 (ThermoFisher #A22287, 1:40 in methanol) for 30 mins at room temperature. Samples were then washed three times in PBS, and mounted on UltraClear microscope slides (Denville Scientific Cat# M1021) using 10 µL ProLong Gold antifade reagent (Invitrogen, #P36930) per coverslip.

#### Fluorescence microscopy

Images were acquired using a Nikon A1r resonant scanning Eclipse Ti2 HD25 confocal microscope with a 10x (Nikon #MRD00105, CFI60 Plan Apochromat Lambda, N.A. 0.45), 20x (Nikon #MRD00205, CFI60 Plan Apochromat Lambda, N.A. 0.75), and 60x (Nikon #MRD01605, CFI60 Plan Apochromat Lambda, N.A. 1.4) objectives, operated by NIS-Elements AR v4.5 acquisition software. Laser intensities and acquisition settings were established for individual channels and applied to entire experiments. Junctional enrichment assay images were collected at 0.07 µm/pixel resolution with 0.07 µm z-step sizes, denoised, and deconvoluted with Richardson-Lucy 3D deconvolution. Image analysis was conducted with Nikon Elements. For the Junction Enrichment analysis, junction enrichment was calculated as a ratio between the corrected mean intensity at junctions (indicated by ZO-1) and the corrected total intensity of the cells in contact. Corrected mean intensities at the regions of interests were obtained by background subtracting mean intensities of the empty vector condition.

#### CELSR signaling assay

Signaling assays were performed as previously described^84,115,116^. HEK293 cells were co–transfected with 0.35 μg CELSR DNA or empty vector, 0.35 μg GloSensor reporter plasmid (Promega, E2301), 9 ng β2 adrenergic receptor, and 2.8 μL of the transfection reagent Fugene 6 (Promega, PRE2693). After a 24–h incubation, the transfected cells were detached and seeded (50,000 cells per well) in a white 96–well assay plate. Following another 24–h incubation, the DMEM was replaced with 100 μL Opti– MEM (Gibco, 31985079) and incubated for 30 min. To each well was then added 1 μL GloSensor substrate and 11 μL FBS. Luminescence measurements were taken after 20 min to allow for equilibration.

### ^107^Statistical Analysis

All statistical analysis was performed using Graphpad Prism 10. For the cell aggregation assays, cell-cell junction assays, and signaling assays, all experiments were performed in at least N=3 independent experiments in at least triplicate. For all assays, the one-way ANOVA with Tukey’s post hoc correction for multiple comparisons was used to assess statistical significance.

## Acknowledgements

We thank Dr. James Fuller, Dr. Minglei Zhao, and Dr. Navid Bavi for valuable discussion and advice regarding cryo-EM data processing and model building. We thank Dr. Zhiqing Wang of the NCEF facility for assistance in cryo-EM data collection. This research was, in part, supported by the National Cancer Institute’s National Cryo-EM Facility at the Frederick National Laboratory for Cancer Research under contract 75N91019D00024. We thank Dr. Maxwell Watkins and Dr. Jesse Hopkins for assistance in SAXS and MALS data collection and processing. This research used resources of the Advanced Photon Source, a U.S. Department of Energy (DOE) Office of Science User Facility operated for the DOE Office of Science by Argonne National Laboratory under Contract No. DE-AC02-06CH11357. BioCAT was supported by grant P30 GM138395 from the National Institute of General Medical Sciences of the National Institutes of Health. The mass photometry analysis was performed by the Biophysics Core in Research Resources Center of University of Illinois at Chicago. Software used in the project was installed and configured by SBGrid. The content is solely the responsibility of the authors and does not necessarily reflect the official views of the National Institute of General Medical Sciences or the National Institutes of Health. This research was supported by the following grants: National Institutes of Health Grant R35GM148412 to D.A., National Institutes of Health Grant F32GM142266 to S.J.B., National Institutes of Health Grant R00MH117235 to R.C.S., and the Alfred P. Sloan Foundation Sloan Research Fellowship to RCS.

## Author contributions

S.J.B., D.A., and R.C.S. conceptualized and designed the study and its methodology. S.J.B., K.G., S.P.K., T.S., E.D., and J.L. conducted the experimental investigations. S.J.B. designed the visualizations in the manuscript with help from S.P.K. and feedback from D.A. D.A. and R.C.S. supervised the work. S.J.B. wrote the original draft of the manuscript. S.J.B. revised and edited the manuscript with feedback from D.A., R.C.S, and S.P.K.

## Competing interests

All authors declare that they have no competing interests.

## Data and materials availability

All data are available in the main text or the supplementary materials. The final model and cryo-EM data have been deposited into the Protein Data Bank under PDB ID 8VY2 and EMDB ID EMD-43644. Small angle X-ray scattering data is available at SASBDB under the accession codes SASDVS7 and SASDVT7. Plasmids and other reagents are available upon reasonable request to the corresponding authors, Demet Araç and Richard C. Sando.

## Notes

### Competing Interest Statement

The authors have declared no competing interest.

### Summary of Updates

Revised Figures and Text in accordance with reviewer comments.

