## Supplemental Information for "Structural basis for regulation of CELSR1 by a compact module in its extracellular region"

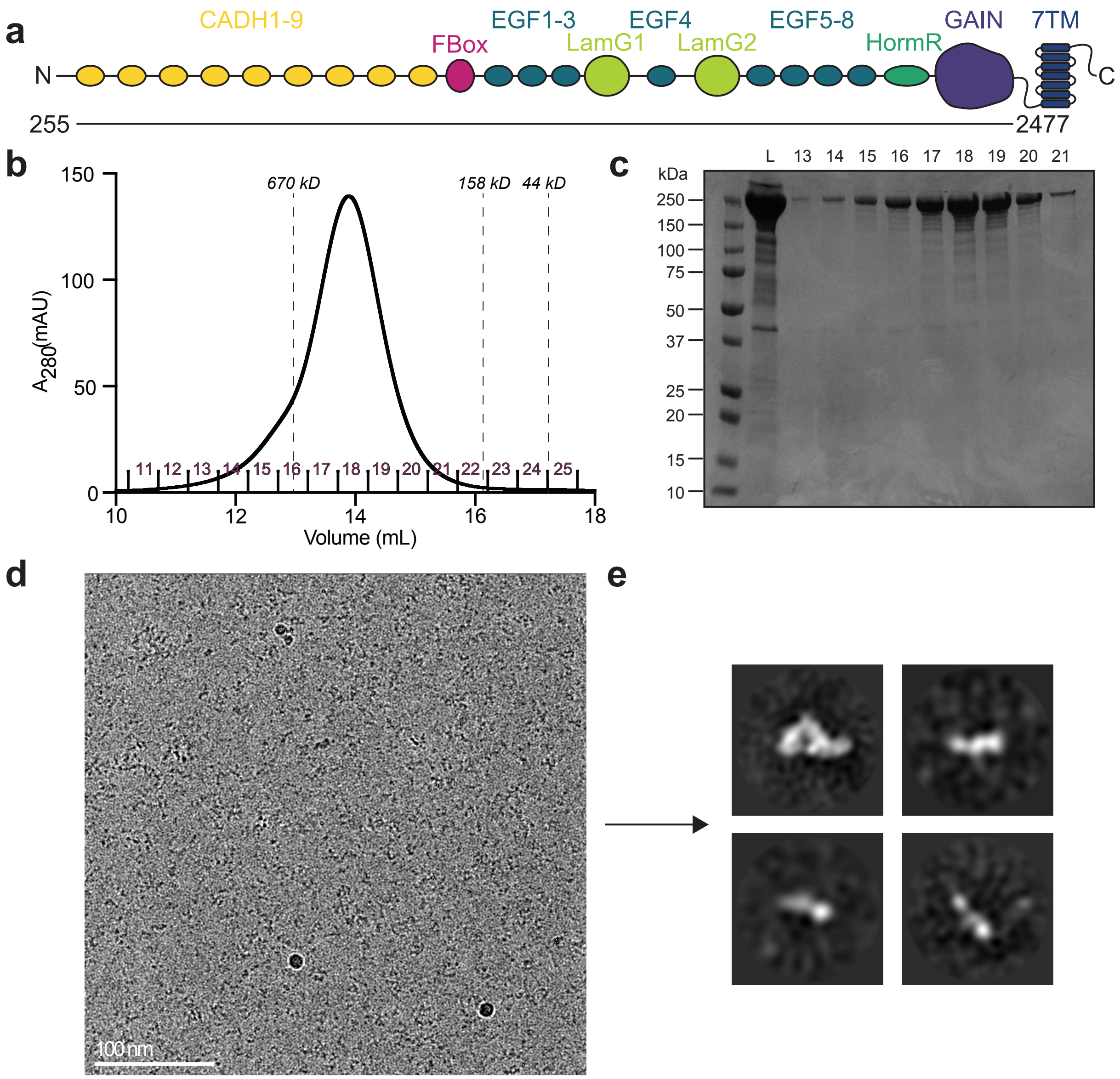


**Supplementary Fig. 1. Purification and cryo-EM screening of the CELSR1 ECR. a** CELSR1 domain diagram with horizontal line showing the extent of the construct used for the structural studies. **b** Size exclusion chromatogram of CELSR1 purification with fraction numbers labeled in purple and size exclusion standards shown as vertical dashed lines. **c** Reducing SDS-PAGE of fractions from SEC trace in **b**. L: fraction loaded onto SEC. **d** Representative screening micrograph of CELSR1 ECR. without added calcium. **e** 2D averages generated from screening micrographs without added calcium.

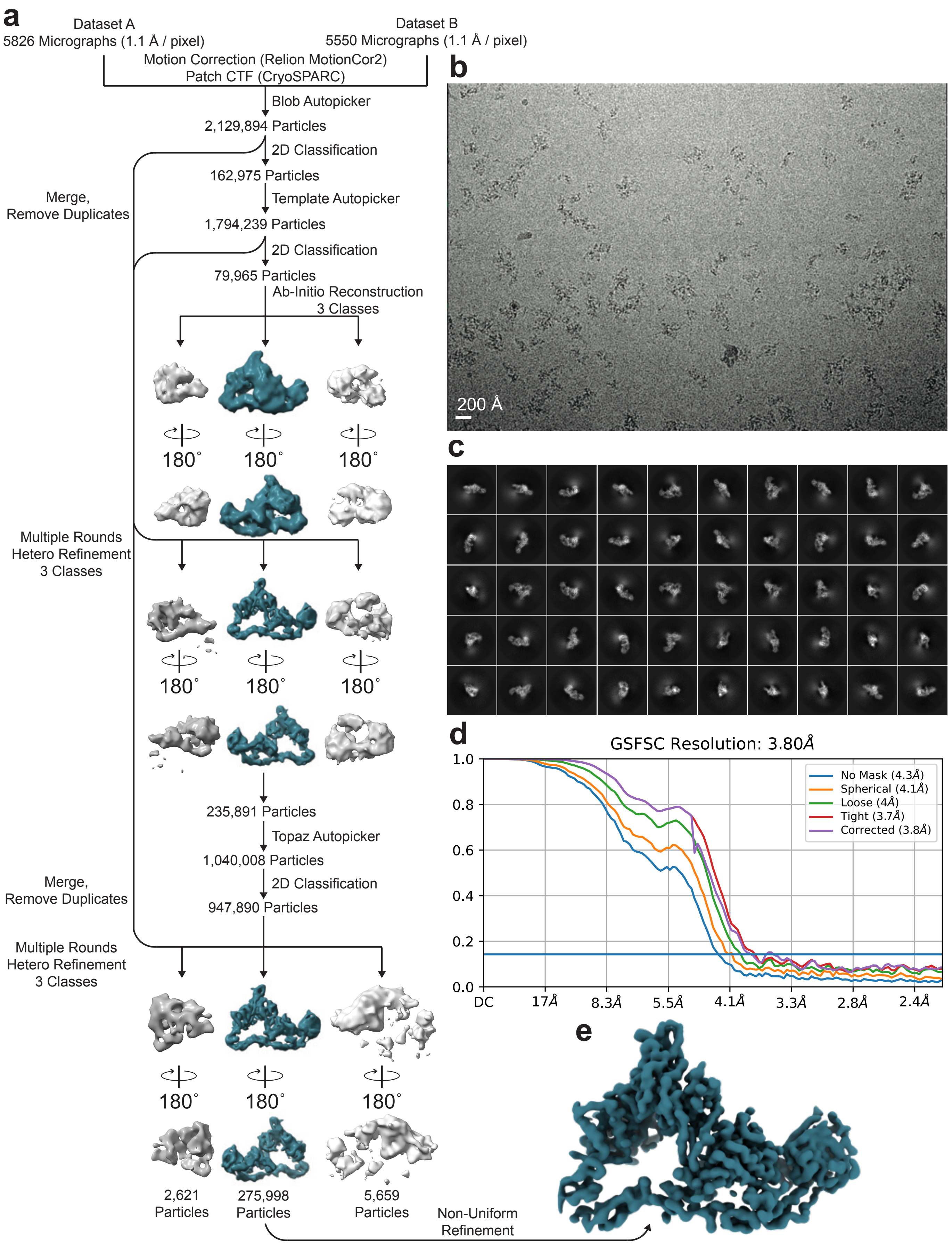

a
Dataset A
5826 Micrographs (1.1 Å / pixel)
Dataset B
5550 Micrographs (1.1 Å / pixel)
Motion Correction (Relion MotionCor2)
Patch CTF (CryoSPARC)
Blob Autopicker
2,129,894 Particles
2D Classification
162,975 Particles
Template Autopicker
1,794,239 Particles
2D Classification
79,965 Particles
Ab-Initio Reconstruction
3 Classes
Merge,
Remove Duplicates
180°
Multiple Rounds
Hetero Refinement
3 Classes
235,891 Particles
Topaz Autopicker
1,040,008 Particles
2D Classification
947,890 Particles
2,621
Particles
275,998
Particles
5,659
Particles
Non-Uniform
Refinement
b
200 Å
c
d
GSFSC Resolution: 3.80Å
No Mask (4.3Å)
Spherical (4.1Å)
Loose (4Å)
Tight (3.7Å)
Corrected (3.8Å)
1.0
0.8
0.6
0.4
0.2
0.0
DC
17Å
8.3Å
5.5Å
4.1Å
3.3Å
2.8Å
2.4Å
e

**Supplementary Fig. 2. Cryo-EM data processing workflow for the CELSR1 ECR reconstruction.** The steps used to process the cryo-EM dataset are shown. **a** Flowchart explaining the data processing steps to generate the CELSR1 reconstruction. **b** Example micrograph collected on Krios TEM. **c** Representative 2D averages of the final particle set. **d** Fourier shell correlation curve showing resolution estimation for the CELSR1 reconstruction. **e** Final density map for CELSR1 reconstruction.

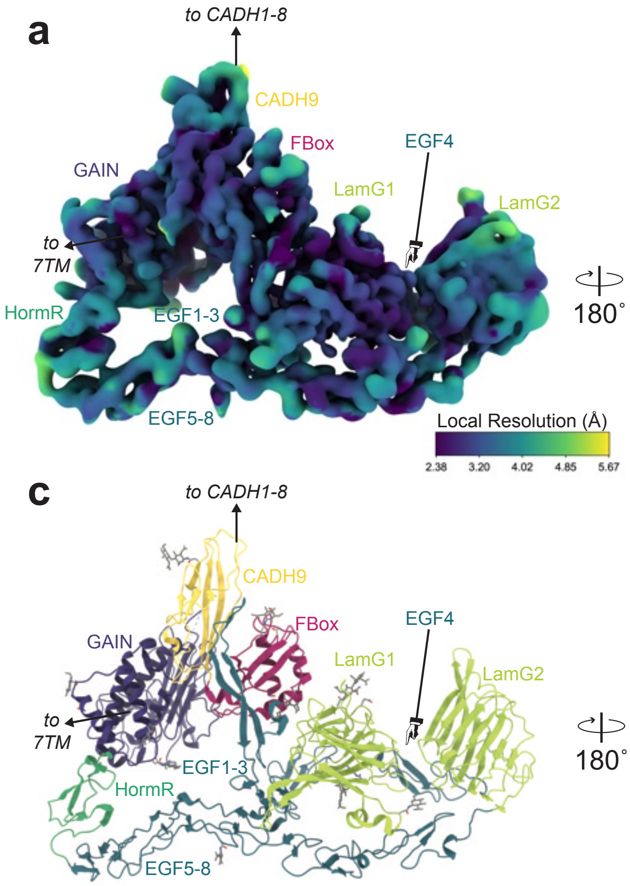


**Supplementary Fig. 3. Local resolution maps of the CELSR1 cryo-EM reconstruction. a,b** Density maps of the CELSR1 reconstruction are shown in two views which are colored by local resolution in Å. Maps were generated using the local resolution tools in cryoSPARC. Domains are labeled and the N- and C-termini are labeled. **c,d** The PDB model for CELSR1 is presented in aligned views to the maps in **a** and **b** for ease of viewing.

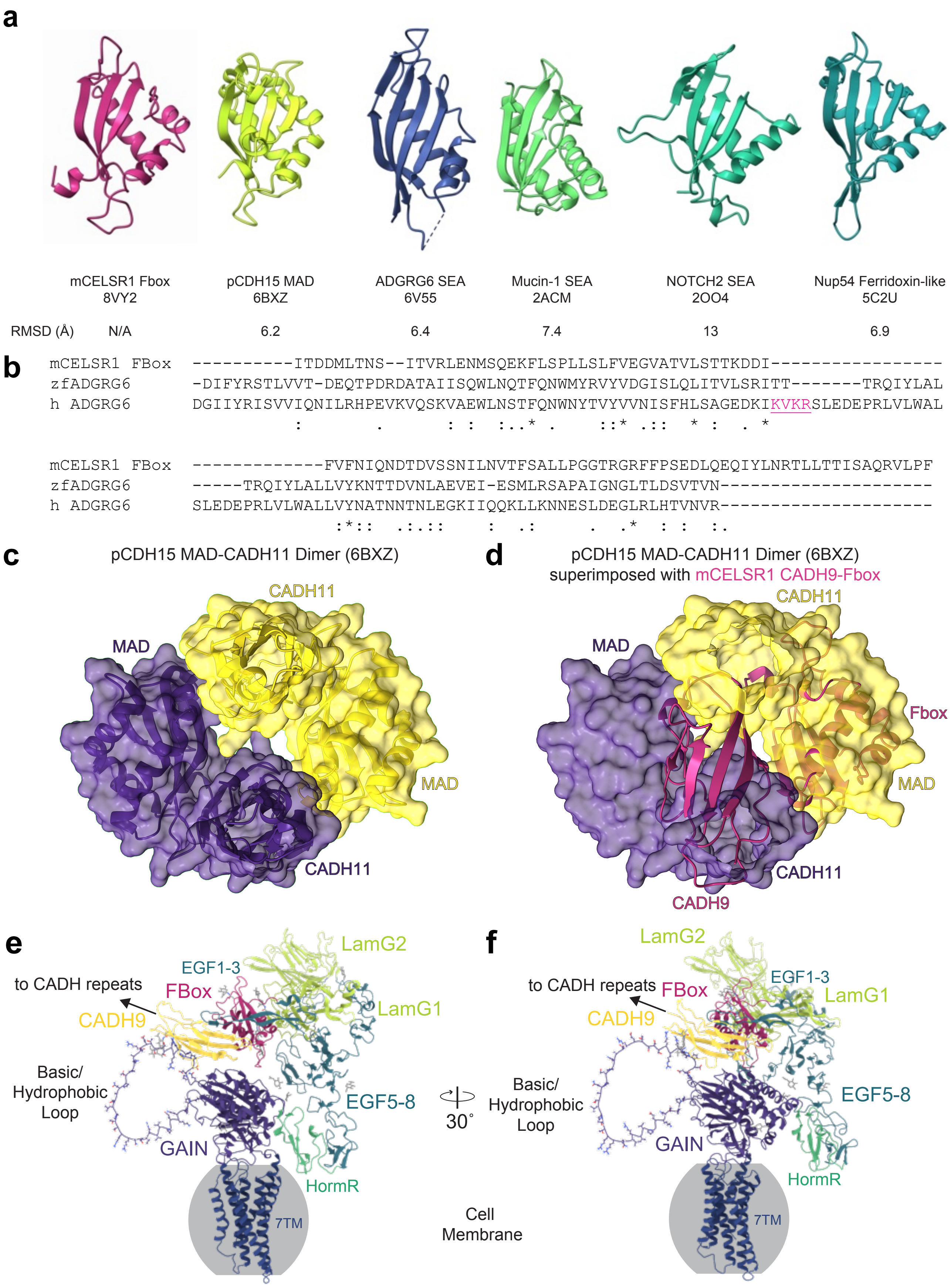

a
mCELSR1 Fbox
8VY2
pCDH15 MAD
6BXZ
ADGRG6 SEA
6V55
Mucin-1 SEA
2ACM
NOTCH2 SEA
2OO4
Nup54 Ferridoxin-like
5C2U
RMSD (Å) N/A 6.2 6.4 7.4 13 6.9
b
mCELSR1 FBox ----------ITDDMLTNS--ITVRLENMSQEKFLSPLLSLFVEGVATVLSTTKDDI-----------------
zfADGRG6 -DIFYRSTLVVT-DEQTPDRDATAIISQWLNQTFQNWMYRVYVDGISLQLITVLSRITT-------TRQIYLAL
h ADGRG6 DGIIYRISVVIQNILRHPEVKVQSKVAEWLNSTFQNWNYTVYVVNISFHLSAGEDKIKVKRSLEDEPRLVLWAL
: . : : :..* . ::* .:: * : . *
mCELSR1 FBox -------------FVFNIQNDTDVSSNILNVTFSALLPGGTRGRFFPSEDLQEQIYLNRTLLTTISAQRVLPF
zfADGRG6 -----TRQIYLALLVYKNTTDVNLAEVEI-ESMLRSAPAIGNGLTLDSVTVN---------------------
h ADGRG6 SLEDEPRLVLWALLVYNATNNTNLEGKIIQQKLLKNNESLDEGLRLHTVNVR---------------------
:*:: .:.:: : .: . .* : : :.
c
pCDH15 MAD-CADH11 Dimer (6BXZ)
CADH11
MAD
MAD
CADH11
d
pCDH15 MAD-CADH11 Dimer (6BXZ)
superimposed with mCELSR1 CADH9-Fbox
CADH11
MAD
Fbox
MAD
CADH11
CADH9
e
LamG2
EGF1-3
to CADH repeats
FBox
LamG1
CADH9
Basic/
Hydrophobic
Loop
EGF5-8
GAIN
HormR
7TM
30˚
Cell
Membrane
f
LamG2
EGF1-3
to CADH repeats
FBox
LamG1
CADH9
Basic/
Hydrophobic
Loop
EGF5-8
GAIN
HormR
7TM

**Supplementary Fig. 4. Analysis of the CELSR1 structure reveals the experimental structure of the Fbox domain and a hypothesis for membrane orientation. a** The experimental CELSR1 Fbox structure is shown in a similar orientation to structural homologs identified using the DALI webserver. The Fbox domain is structurally similar to MAD, SEA, and Ferridoxin-like domains. The all-atom RMSD is shown below. **b** Sequence alignments of Fbox domain with zebrafish and human ADGRG6 reveal that the CELSR1 Fbox domain does not have a furin cleavage site where some other SEA domains do. The furin cleavage site in human ADGRG6 is shown in magenta and is underlined. **c** The pCDH15 CADH11-MAD dimerization site is shown and **d** the CADH9-Fbox orientation is shown to sterically slash with the pCDH15 mode of dimerization. **e,f** AlphaFold2 predictions for the GAIN-7TM region of CELSR1 overlaid with the CADH9-GAIN CMM suggests a possible membrane orientation for the CADH9-GAIN CMM.

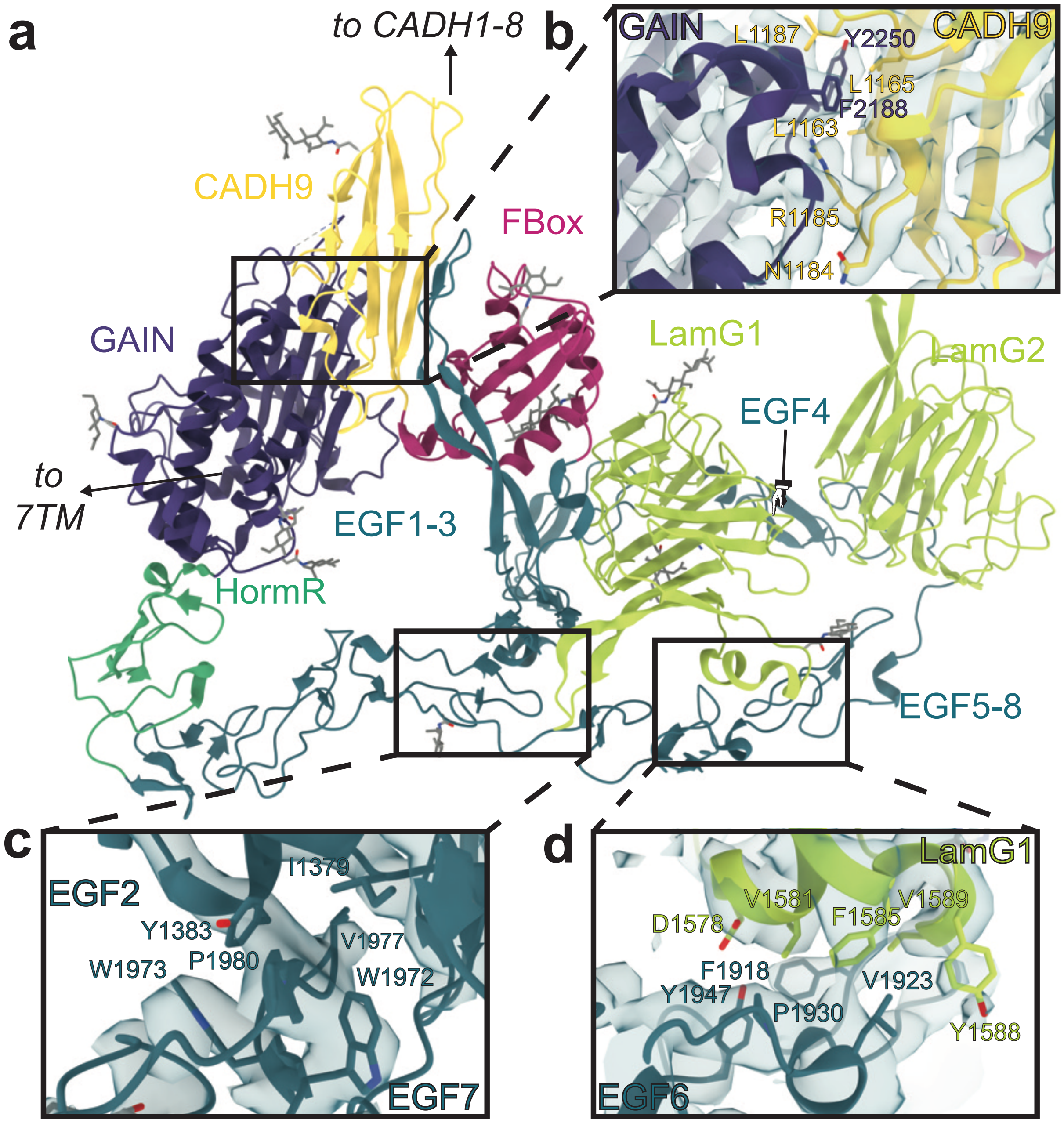


**Supplementary Fig. 5. Interdomain interfaces stabilizing the CELSR1 ECR compact bundle rendered using ChimeraX, with density shown. a** Cartoon representation of CELSR1 ECR structure with key interfaces between **b** CADH9 and GAIN domains, **c** EGF2 and EGF7, and **d** EGF6 and LamG1 with key side chains shown in sticks and density shown using blue surface representation.

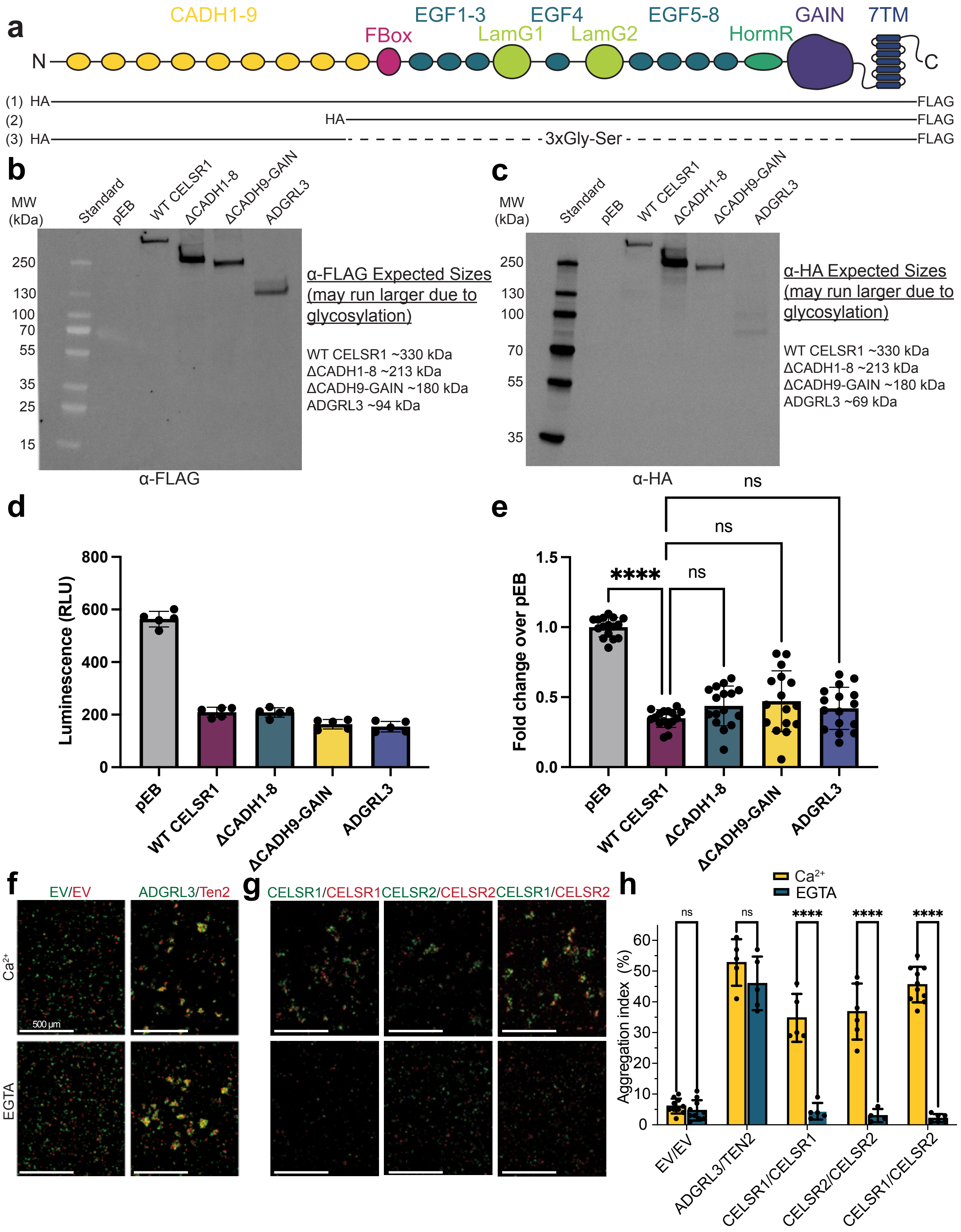


**Supplementary Fig. 6. Functional analysis of CELSR ECR constructs. a** Domain boundaries of deletion constructs used in this study. Black solid lines show domain boundaries with dotted lines indicating regions that are connected with linkers. **b** Experimental constructs were probed using western blotting against the C-terminal FLAG tag or **c** N-terminal HA tag to confirm the construct size. Expected sizes are noted, some constructs may run larger than expected due to glycosylation. **d** Signaling assay for CELSR function. Experimental constructs are co-transfected with $\beta_2$AR and CELSR1 constructs result in lower cAMP relative to empty vector control. ADGRL3 is used as a positive control as previously described[1–3]. Representative raw experiment shown. **e** Compiled results of N=3 independent biological experiments performed at least in triplicate presented as fold change over empty vector. **f,g** Representative cell aggregation assay images showing effect of EGTA on cell aggregation mediated by ADGRL3/Ten2 and CELSR isoforms. EGTA does not affect ADGRL3/Ten2-mediated aggregation but disrupts CELSR-mediated aggregation. CELSR1/CELSR2 can aggregate heterophilically. **h** Quantification of **f,g** as aggregation index. one-way ANOVA with Tukey's correction for multiple comparisons was performed to assess statistical significance between CELSR constructs. *** corresponds to $p = 0.0001$, **** corresponds to $p < 0.0001$. Each construct was compared to each other construct for the statistical testing, but only certain comparisons are shown for clarity.

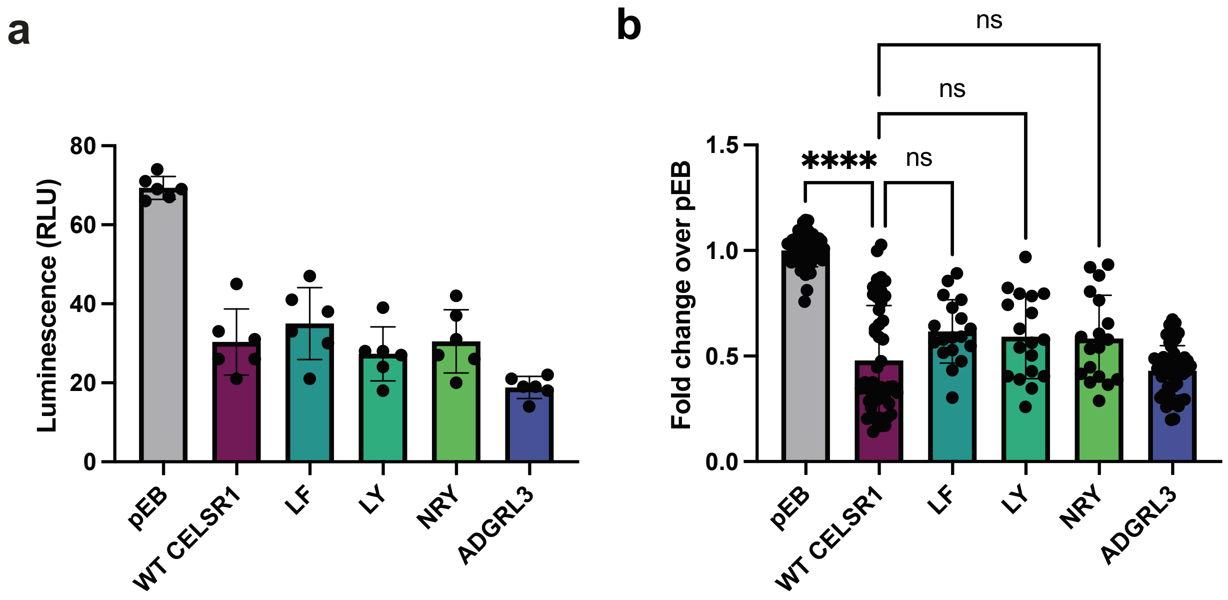


**Supplementary Fig. 7. Functional analysis of CADH9/GAIN interfacial point mutations.** Experimental constructs are co-transfected with β2AR and CELSR1 constructs result in lower cAMP relative to empty vector control. LF, L1163A/F2188A; LY, L1163A/Y2250A; NRY, N1184A/R1185A/Y2250A. ADGRL3 is used as a positive control as previously described[1–3]. **a** Representative raw experiment shown. **b** Compiled results of N=3 independent biological experiments performed at least in triplicate presented as fold change over empty vector. one-way ANOVA with Tukey's correction for multiple comparisons was performed to assess statistical significance between CELSR constructs. *** corresponds to $p = 0.0001$, **** corresponds to $p < 0.0001$. Each construct was compared to each other construct for the statistical testing, but only certain comparisons are shown for clarity.

| Residue Name | Buried Surface Area (Å$^2$) | Contacting residues on opposing domain (Residues <6 Å in distance) |
|---|---|---|
| | | **CADH9/GAIN interface** |
| Q1159 | 16.6 | P2348 |
| G1160 | 8.1 | Y2250 |
| E1162 | 31.0 | Y2250 |
| L1163 | 99.7 | G2187, F2188, R2248, T2249 |
| L1165 | 31.3 | F2188, R2248, Y2250 |
| D1180 | 40.4 | F2188, T2193 |
| D1182 | 3.7 | T2193, R2194 |
| N1183 | 42.5 | F2188, T2193, R2194, E2195 |
| N1184 | 53.6 | R2194, E2195, A2196 |
| R1185 | 121.9 | F2188, R2194, E2195, A2196, T2249, Y2250, L2251, R2252 |
| P1186 | 32.4 | R2252, Y2250 |
| L1187 | 34.5 | T2249, Y2250, L2251, R2252 |
| E1188 | 16.3 | Y2250, R2252 |
| A1189 | 15.2 | Y2250 |
| M1191 | 7.5 | Y2250 |
| G2187 | 12.5 | L1163 |
| F2188 | 107.2 | L1163, S1164, L1165, D1180, L1181, N1183, R1185 |
| T2193 | 41.7 | D1180, D1182, N1183, R1185 |
| R2194 | 30.2 | D1182, N1183, N1184, R1185 |
| E2195 | 26.8 | N1183, N1184, R1185 |
| A2196 | 31.5 | N1184, R1185 |
| R2248 | 55.7 | E1162, L1163, L1165 |
| T2249 | 24.6 | L1163, R1185, L1187 |
| Y2250 | 110.7 | G1160, E1162, L1165, R1185, P1186, L1187, E1188, A1189, M1191 |
| L2251 | 22.4 | R1185, L1187 |
| R2252 | 73.0 | R1185, L1187, E1188 |
| P2348 | 14.9 | Q1159 |
| | | **EGF2/EGF7 interface** |
| T1371 | 23.2 | W1972 |
| I1379 | 31.4 | W1972, C1978, G1979, P1980 |
| D1380 | 37.7 | V1977, C1978, P1980 |
| Y1383 | 123.7 | V1977, C1978, P1980, W1973, G1974 |
| S1384 | 7.2 | W1973 |
| N1385 | 16.3 | W1973, N1975 |
| W1972 | 44.1 | T1371, I1379 |
| W1973 | 47.3 | Y1383, S1384, N1385 |
| G1974 | 4.2 | Y1383 |
| N1975 | 4.7 | N1385 |
| V1977 | 57.3 | D1380, Y1383 |
| C1978 | 9.7 | I1379, D1380, Y1383 |
| G1979 | 15.1 | I1379, D1380, Y1383 |
| P1980 | 42.5 | I1379, Y1383 |
| | | **LamG1/EGF6 interface** |
| D1578 | 41.2 | F1918, R1945, Y1947, Y1959 |
| A1580 | 33.3 | P1930, Y1959 |
| V1581 | 49.4 | F1918, A1925, P1930, Y1947 |
| H1584 | 32.8 | K1932 |
| F1585 | 54.3 | A1925, L1928, P1930 |
| Y1588 | 28.2 | K1921, L1928 |
| V1589 | 76.8 | F1918, G1919, K1920, K1921, V1923, L1928 |
| G1590 | 12.0 | G1919, K1920, K1921 |
| N1591 | 19.7 | G1919, K1920 |
| Y1592 | 34.3 | F1918, G1919 |
| F1918 | 67.0 | D1578, V1581, V1589, Y1592 |
| G1919 | 42.0 | V1589, G1590, N1591, Y1592 |
| K1920 | 28.8 | V1589, G1590, N1591 |
| K1921 | 25.4 | Y1588, V1589, G1590 |
| V1923 | 8.8 | V1589 |
| A1925 | 20.0 | V1581, F1585 |
| L1928 | 71.3 | F1585, Y1588, V1589 |
| P1930 | 38.6 | A1580, V1581, F1585 |
| K1932 | 32.2 | H1584 |
| R1945 | 6.7 | D1578 |
| Y1947 | 25.5 | D1578, V1581 |
| Y1959 | 26.6 | D1578, A1580 |

**Supplementary Table 1: List of interactions in the CELSR1 CADH9-GAIN structure.**

### Supplementary Methods

#### Western blotting

Western blotting was performed as previously described[4,5]. Briefly, HEK293T cells were transfected with 2 micrograms of CELSR constructs using LipoD293 (SignaGen Laboratories SL100668). 48 hours later, cells were washed and resuspended in 500 µL of solubilization buffer (20 mM HEPES pH 7.4, 150 mM NaCl, 2 mM MgCl2 0.1mM EDTA, 2 mM CaCl2, 1% (v/v) Triton X-100). After clarification by high-speed centrifugation, cell lysates were run on SDS-PAGE and transferred to nitrocellulose using a wet transfer system. After blocking, membranes were incubated with either anti-FLAG or anti-HA antibody conjugated to iFluor 488 or iFluor 647, respectively. The next day, membranes were washed and imaged at respective wavelengths using the ChemiDoc imaging system.
